# Directed evolution of L-DOPA sensing and production enables efficient incorporation of an expanded genetic code into proteins

**DOI:** 10.1101/2025.06.26.661824

**Authors:** Andrew R. Gilmour, Qiyao Wei, Chad Wang, Jessica Hellinger, Prithvijeet Singh Kharnal, Vidya Pandarinath, Simon d’Oelsnitz, Eli Powell, Jennifer S. Brodbelt, Andrew D. Ellington, Ross Thyer

**Affiliations:** Systems, Synthetic, and Physical Biology, Rice University, Houston, TX; Department of Bioengineering, Rice University, Houston, TX; Department of Molecular Biosciences, The University of Texas at Austin, Austin, TX; Department of Chemistry, The University of Texas at Austin, Austin, TX; Department of Integrative Biology, The University of Texas at Austin, Austin, TX; Department of Chemical and Biomolecular Engineering, Rice University, Houston, TX

## Abstract

L-3,4-dihydroxyphenylalanine (L-DOPA), is a versatile catechol-containing amino acid derived from L-tyrosine that is both a key metabolite for the production of many valuable natural products and a naturally occurring post-translational modification in proteins. While advances in microbial natural product biosynthesis and site-specific incorporation of L-DOPA have both paved the way for relevant applications of L-DOPA, progress has been stifled by a lack of robust *in vivo* detection tools and high-efficiency biosynthetic enzymes. Here, we report and engineer a LysR-family transcription factor, PP_2251, that can serve as a transcriptional regulator for L-DOPA with a dynamic range of >100-fold when coupled with a fluorescent reporter. Leveraging this improved regulator in conjunction with compartmentalized partnered replication (CPR), we engineered a promiscuous *E. coli* flavin-dependent monooxygenase (HpaB) for enhanced L-DOPA biosynthesis. This represents the first use of high-throughput emulsion PCR to improve a metabolic enzyme and yielded HpaB variants that showed >40-fold activity within our genetic circuit and a 6-fold increase in L-DOPA titer compared to wild-type HpaB. We further demonstrated that the highest performing HpaB variant was capable of supporting efficient translational incorporation of L-DOPA into protein, achieving 97% incorporation efficiency and yields approaching 250 mg.L^-1^. Collectively, these advancements enable the *in vivo* detection, biosynthesis, and site-specific incorporation of L-DOPA for synthetic and chemical biology applications.

## Main

There is broad interest in microbial L-DOPA biosynthesis as the first committed (and rate-limiting) step towards the production of many biomedically important natural products, including benzylisoquinoline alkaloids, a diverse family of plant-derived compounds including opiate analgesics, and the catecholamine neurotransmitters, dopamine and epinephrine^1,2^. However, efforts to engineer these biosynthetic pathways have been complicated by the reactivity of L-DOPA imposing a burden on cell fitness as well as a lack of quantitative high-throughput screening approaches to parse large enzyme libraries. Prior work has relied on abiotic or enzymatic oxidation of L-DOPA to melanin or betalain pigments^3–5^. However, these screens do not directly report L-DOPA concentrations, and each suffers from fundamental limitations that affect their dynamic range: melanin formation is toxic while *E. coli* does not retain betalains inside the cell, instead secreting them into the culture medium^6^.

In contrast, transcriptional regulators are powerful tools for guiding the improvement of biosynthetic enzymes and enable large gene libraries to be interfaced with a variety of genetic outputs to facilitate high-throughput screening^7^. Genetic circuits and screening approaches can be further tailored to balance the burdens on host fitness with increases in enzyme activity to ensure cryptic selection pressures are not confounding the outcomes. These include emulsion-PCR techniques such as compartmentalized partnered replication (CPR) whereby improved gene variants are coupled to the expression of a thermostable DNA polymerase. This approach decouples host fitness from enzyme activity and confers strong selective pressure by exponential amplification of improved gene sequences^8,9^. In addition, using the tools of synthetic biology, CPR gene circuits can easily be designed to enable counter-selections with equally powerful enrichment of desirable phenotypes.

Beyond its applications in biosynthesis, L-DOPA as an amino acid in proteins plays a crucial role in the properties of many biomaterials, including adhesive proteins produced by marine invertebrates and the durable biopolymers in squid beaks^10–12^. Its occurrence is primarily the result of the post-translational modification of tyrosine by the enzyme tyrosinase, although it can also be introduced into proteins via misincorporation in place of tyrosine when L-DOPA concentrations are unnaturally elevated (typically when L-DOPA is being used therapeutically to manage neurodegenerative disorders^13^). However, there is no natural defined encoding of L-DOPA into proteins, and its occurrence is extremely heterogenous, driven by random misincorporation, susceptibility to oxidative damage, or enzymatic modification. This heterogeneity limits the use of L-DOPA and its unique catechol functionality for engineering proteins with consistent biophysical or enzymatic properties, including bioconjugation, metalloprotein engineering, and radical catalysis^14^. One solution to this problem is the use of orthogonal translational machinery to site-specifically introduce L-DOPA during translation. Unfortunately, this approach requires supplementation with exogenous L-DOPA, a significant barrier for production at scale.

Concurrent biosynthesis and incorporation of L-DOPA into proteins avoids the need for supplementation and offers a tractable route to low-cost production. There are three canonical routes for L-DOPA biosynthesis: copper-containing tyrosinase, non-heme tyrosine hydroxylases, and the heme-dependent tyrosine hydroxylase CYP76AD1; however, each has limitations for use in bacteria^15–17^. Tyrosinases will act indiscriminately on tyrosine residues in proteins and the enzymes are prone to further oxidize L-DOPA to reactive quinone species^18^. The heme-dependent tyrosine hydroxylase cytochrome P450 CYP76AD1 can also further act on L-DOPA, converting it into cyclo-DOPA. Finally, non-heme tyrosine hydroxylases require the metabolically expensive cofactor tetrahydromonapterin and a dedicated two-enzyme cofactor recycling pathway in *E. coli*^16^.

Non-native biosynthetic pathways for L-DOPA also exist. Tyrosine phenol-lyase (TPL) can be used to directly biosynthesize L-DOPA from alternate building blocks and support incorporation into proteins, but this pathway is difficult to interface with central metabolism due to the requirement for free catechol which is not a standard metabolite in *E. coli*^19^. Another option is 4-hydroxyphenylacetate 3-monooxygenase (HpaB), a flavin-dependent monooxygenase with high substrate promiscuity towards phenolic compounds, including weak activity on L-tyrosine^20,21^. In contrast to the other alternatives, HpaB is natively expressed in some *E. coli* strains and has simple cofactor requirements, needing only FADH_2_ and a flavin reductase (HpaC) (**Figure 1a**). Furthermore, there is an abundance of high-resolution structural data available for both the *E. coli* enzyme and the structurally homologous ortholog from *Thermus thermophilus* to guide engineering efforts^22–24^. Here we report the isolation and engineering of a transcriptional regulator for L-DOPA, along with the use of ultra-high-throughput emulsion-PCR methods to engineer *Ec*HpaB variants optimized for L-DOPA biosynthesis and show that an improved biosynthetic pathway supports higher incorporation of L-DOPA into recombinant proteins.

**Figure 1:**
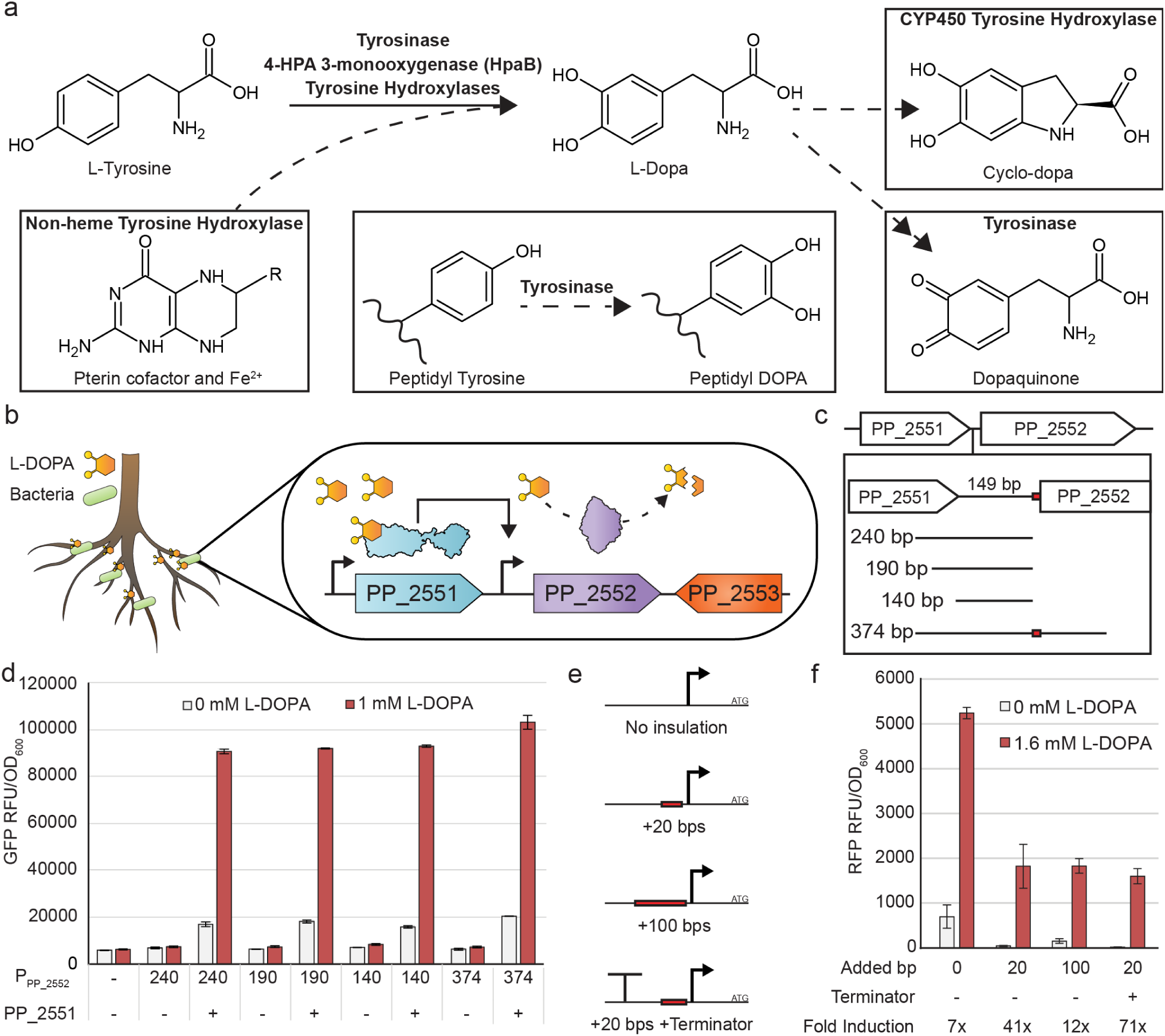
Characterization and optimization of the PP_2552 promoter. a) Diagram of biosynthetic pathways for generating L-DOPA. In boxes are the undesirable reactions or co-factor requirements that limit the use of the specified enzyme for use in producing and sensing L-DOPA. b) Schematic for *Pseudomonas putida* KT2440 and its operon containing a LysR-family transcriptional regulator (PP_2551), a L-DOPA decarboxylase (PP_2552), and a major facilitator superfamily transporter (PP_2553). c) Design of the four different putative promoters based on the intergenic region between PP_2551 and PP_2552. The red mark indicates the native RBS which is replaced with a strong RBS when not included. d) Fluorescence assay of the four putative promoters using a sfGFP reporter protein with and without exogenous L-DOPA and expression of PP_2551. Data represents the average fluorescence of three independent clones normalized to cell density ± s.e.m. e) Schematic of progressive modifications to the upstream region of the PP_2552 promoter to provide insulation from the surrounding plasmid context. f) Screening of insulated promoter variants using an mScarlet-I reporter. Data represents the average fluorescence of three biological replicates normalized to cell density ± std.

## Results

### PP_2551: a L-DOPA responsive transcription regulator

There is currently no known transcriptional regulator which natively responds to L-DOPA. To identify a transcription regulator for L-DOPA we pursued three separate strategies: (i) evaluating known transcriptional regulators with native ligands that share structural similarity with L-DOPA, (ii) screening highly promiscuous multidrug sensors that may possess basal recognition of L-DOPA, and (iii) investigating uncharacterized controllers of L-DOPA-specific enzymes (**Figure S1**). For the first approach, transcription repressors PcaU (3,4-dihydroxybenzoic acid or 3,4-DHBA)^25^, HpaR (3,4-dihydroxyphenylacetic acid or 3,4-DHPA)^26^, or MidR (L-mimosine)^27^ were co-expressed alongside a green fluorescent protein (GFP) reporter under the control of their cognate promoters, P*_pcaI_*, P*_hpaA_*, and P*_midA_* respectively. However, none of these regulators were observed to induce GFP expression when induced with 1 mM L-DOPA. Two promiscuous regulators that control multidrug efflux pumps (QacR and TtgR) were similarly evaluated using the P*_qacA_* and P*_ttgA_* promoters respectively^28,29^, but neither showed any response towards L-DOPA.

Failure of these initial efforts motivated us to investigate the regulatory architecture of pathways that make or act on DOPA and potentially identify uncharacterized DOPA-modulated transcription regulators. DOPA decarboxylases have been identified in environmental bacteria^30,31^, and a L-DOPA decarboxylase from the plant-associated *Pseudomonas putida* KT2440 (PP_2552) that was reported to have high selectivity for L-DOPA is located adjacent to a LysR-family transcription regulator (PP_2551) and a major facilitator superfamily transporter (PP_2553) (**Figure 1b**)^31^. Since transcription of the gene cluster encoding PP_2552 was observed in the presence of L-DOPA, but not L-tyrosine, even at relatively high concentrations (5 mM of L-Tyr), we hypothesized that PP_2551 likely controlled the expression of PP_2552 in a L-DOPA-specific manner.

To investigate DOPA-mediated regulation by PP_2251 in *E. coli*, we initially sought to identify the minimal promoter sequence for PP_2552. Given that no promoter or operator sites could be identified within the intergenic region between PP_2551 and PP_2552, four truncations of the surrounding region were constructed and placed upstream of a sfGFP fluorescent reporter (**Figure 1c**). In three designs, the nine base pairs immediately upstream of the PP_2552 start codon were replaced with a strong *E. coli* translation initiation region. Each fragment was evaluated for promoter activity in the presence and absence of both PP_2551 and exogenous L-DOPA in *E. coli* DH10B (**Figure 1d**). All four fragments demonstrated promoter activity that was dependent upon the presence of both the transcription regulator and L-DOPA. The smallest (140 bp) appeared to contain all the required regulatory elements and was adopted for further optimization. To buffer the promoter against transcriptional interference from the surrounding genetic context of the plasmid, two different insulating elements were evaluated (**Figure 1e-f**). The combined addition of both 20 bp of neutral DNA context (taken from the original characterization plasmid, but not present in *Pseudomonas*) and the rrnBt transcriptional terminator from Mutalik *et al*., eliminated most background activity and resulted in 71-fold induction when saturated with 1.6 mM L-DOPA, a 10-fold improvement over the original promoter sequence^32^.

### Engineering the PP_2551 transcriptional regulator

PP_2551 requires a high concentration of L-DOPA (5 mM) in *Pseudomonas putida* to maximally induce *PP_2552*^31^, and this lack of sensitivity limits further pathway engineering. An emulsion-based method, Compartmentalized Partnered Replication (CPR), has previously been used to improve the effector specificities of transcription factors, including their ability to distinguish structurally similar ligands (anticipated to be a problem with L-DOPA and L-tyrosine), and was adapted to engineer PP_2551 variants with higher dynamic ranges (**Figure 2a**)^33^. In CPR, variants capable of improved transcriptional activation at increasingly lower L-DOPA concentrations generate a thermostable DNA polymerase that in turn leads to self-amplification of the variant gene during emulsion PCR. To eliminate variants that might result in either a constitutively active phenotype or enable the recognition of L-tyrosine or other cryptic effectors, an additional counter-selection was constructed in which unwanted variants activate expression of the lambda cI repressor that blocks expression of the DNA polymerase, thus preventing amplification during emulsion PCR.

**Figure 2:**
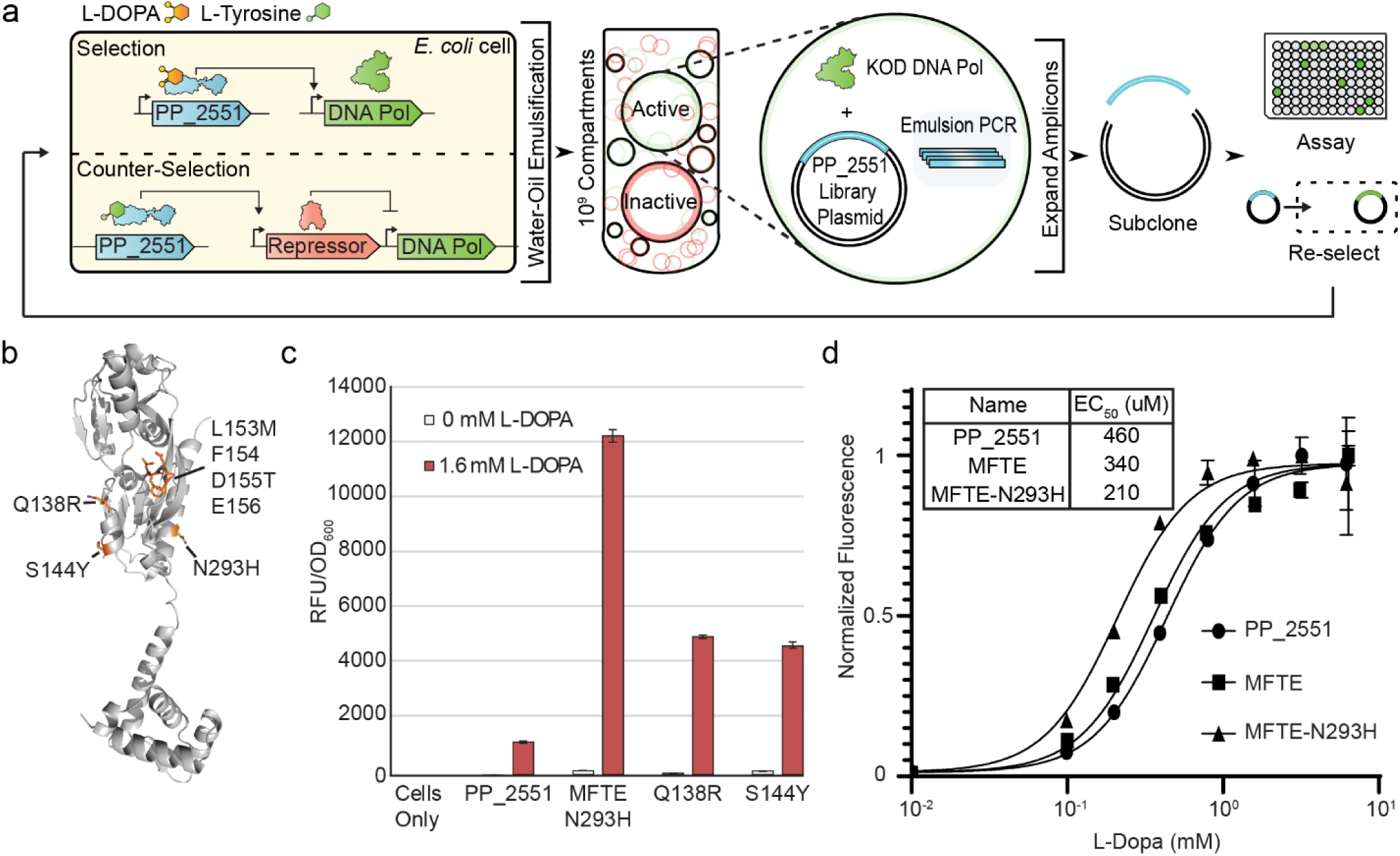
Evolution of the PP_2551 transcriptional regulator for improved response towards L-DOPA. a) Schematic of the Compartmentalized Partnered Replication (CPR) selection workflow for the PP_2551 transcriptional regulator. Variants with greater response to L-DOPA result in increased expression of the KOD DNA Polymerase and subsequently greater enrichment during emulsion PCR. b) AlphaFold structure of PP_2551 with best performing mutations labeled. c) Fluorescence assay of best performing PP_2551 mutations isolated from CPR selections. Data represents the average fluorescence of three biological replicates normalized to cell density ± std. d) EC_50_ determination of PP_2551 wildtype, MFTE, and MFTE-N293H. Data represents the average fluorescence of three biological replicates normalized to cell density ± std

In order to generate libraries for selection, we first generated a homology model with SWISS-MODEL, starting from the putative LysR regulator PA_0477 from *Pseudomonas aeruginosa* (PDB: 2ESN)^34^. Two regions flanking the likely ligand binding cavity were selected for site-saturation mutagenesis: A136/G137/Q138/Y139 and L153/F154/D155/E156 (**Figure 2b**). Following transformation, each library exceeded a minimum of 10-fold coverage; they were then subjected to three rounds of positive selection interspersed with two rounds of counter-selection. The first library yielded only one variant with improved properties, Q138R, while the second resulted in several different mutations, including several off-target mutations (**Figure S2**). The single best variant L153M/D155T/N293H, designated MFTE-N293H, displayed improved induction in response to L-DOPA with no increase in background (**Figure 2c**).

We noticed that our top variant harbored a mutation outside the targeted ligand binding domain (N293H) and hypothesized that it may contribute to improved L-DOPA sensitivity. Therefore, we removed the off-target mutation to generate a variant herein named ‘MFTE’. Titration of PP_2551, MFTE, and MFTE-N293H with a gradient of exogenous L-DOPA concentrations indicated that N293H was in fact functional and important in increasing the sensitivity towards L-DOPA. Ultimately, MFTE alone reduced the EC_50_ from 460 μM to 340 μM, and the addition of N293H further reduced it to 210 μM (**Figure 2d**). These results indicate that methods such as CPR that rely both on library generation and mutations accumulated during amplification can efficiently explore sequence space for functional variants. We also investigated the effect of rich and minimal media compositions and found that rich media degraded the performance of the transcriptional regulator, possibly due to the presence of low levels of L-DOPA resulting from trace oxidation of L-tyrosine which is present at much higher concentrations (**Figure S3a-b**). Given that the same effect would have greatly decreased the already low dynamic range of PP_2551, improving the effector sensitivity of the transcription factor was essential for further engineering of the pathway.

### Using the PP_2551 transcriptional regulator to improve *Ec*HpaB for L-DOPA biosynthesis

To integrate transcriptional regulator development with pathway engineering, we constructed a dedicated *Ec*HpaB expression plasmid which placed *hpaB* under the control of P_LtetO_ and constitutively expressed *hpaC*. A small site-saturation library was constructed around Gly 295 and Leu 411, two positions previously identified by Fordjour *et al*. that had yielded mutants with enhanced L-DOPA production^35^. The library was transformed into cells harboring a fluorescent reporter regulated by the MFTE-N293H PP_2551 variant and 24 highly fluorescent transformants were selected for characterization (**Figure S4a-b**). Multiple mutations were recovered at G295 and L411, but not the previously reported G295A, G295R, or L411M. This was not completely surprising, as previous attempts at optimization had relied on random mutagenesis, rather than site-saturation. Four of the most abundant mutants were reconstructed and re-phenotyped using a fluorescence assay which confirmed they outperformed the parental *Ec*HpaB enzyme (**Figure S4c**), and that the evolved transcriptional regulator was suitably sensitive to L-DOPA.

Having validated the integration of transcriptional regulator with biosynthesis, we modified our previous CPR circuit by co-locating the regulator and DNA polymerase onto the selection plasmid. Since *Ec*HpaB, expressed from the library plasmid, was the sole source of L-DOPA (**Figure 3a**), variants can induce expression of the thermostable DNA polymerase and subsequently be amplified in emulsions relative to their activity. Based on analysis of the *Ec*HpaB structural data, we identified a region flanking the active site that we hypothesized would be a good candidate for site-saturation mutagenesis, consisting of residues 209-212 of the highly flexible β32-β33 loop that undergoes conformational changes upon substrate and cofactor binding in the *Thermus thermophilus* ortholog (*Tt*HpaB)^23^. Following three rounds of CPR selection, amplicons were subcloned and transformed into cells containing a fluorescent reporter plasmid for screening. The first library (G209/S210/A211/Q212) generated many variants with improved activity (**Figure S5**). Gly 209 was largely invariant, with a single clone encoding a proline at that position. In contrast, all variants replaced Ala 211 with arginine except for a single clone encoding a lysine. To confirm that our selection scheme was isolating variants that biosynthesized L-DOPA, culture media from *E. coli* cells expressing variant GMRQ (the first identified) was analyzed using HPLC with an electrochemical detector (HPLC-ECD). A peak consistent with the desired product was found (**Figure S6**).

**Figure 3:**
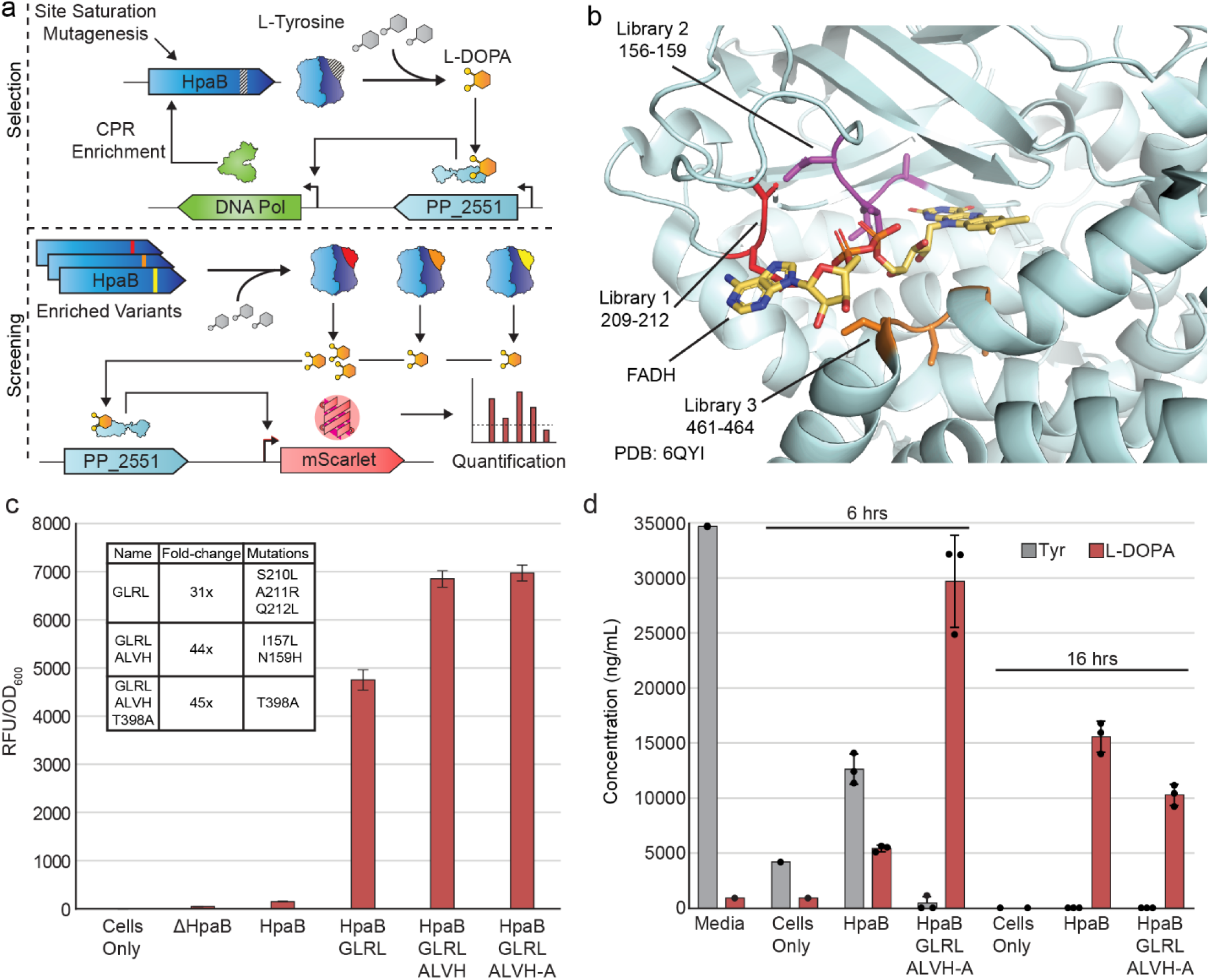
Transcriptional regulator guided evolution of a L-DOPA biosynthesis enzyme, 4-hydroxyphenylacetate 3-monooxygenase (HpaB). a) Schematic of the CPR selection method for HpaB. HpaB mutations that result in increased biosynthesis of L-DOPA are linked to expression of the KOD DNA Polymerase via PP_2551 L-DOPA transcriptional regulator. b) Structural representation of the HpaB active site showing the three site saturation libraries. c) Fluorescence assay of the highest performing HpaB mutations after each round of library selection. Library 3 recovered no mutations from the library region. d) LC-MS quantification of L-DOPA and L-Tyrosine at 6 h and 16 h post-induction of HpaB expression.

Starting from one of the most active variants from the first round (GDRK-R94H), two additional site-saturation libraries were constructed, spanning residues 156-159 in the loop at the beginning of the central beta-sheet domain adjacent to His 155 which forms part of hydrogen bonding network with the substrate phenolic oxygen (A156/I157/V158/N159), and residues 461-464 which form a short linker on the opposite face of the substrate binding cavity (Y461/S462/G463/S464; **Figure 3b**). These libraries were again selected using CPR. Variants with similar sets of conservative mutations were isolated from the second library, while no mutations within the mutagenized region of the third library were recovered. Given that many mutants can be highly context dependent, we selected a panel of the most abundant sequences isolated from the second library and screened them for improved activity with another highly active variant from the first library selection. Four different sequences were grafted into HpaB variant GLRL, and screening revealed the GLRL-ALVH combination greatly improved activity (**Figure S7**). In addition, several off-target mutations that had arisen during selection were also inserted into these variants (**Figure S8**). The addition of T398A ultimately yielded two variants with almost identical activity, GDRK-R94H-SLVN-A and GLRL-ALVH-A (**Figure 3c, S9**). These results emphasize that CPR can be used to broadly redefine the active site for improved activity, putting the protein on a new evolutionary trajectory, including the acquisition of point mutations that further optimize the new architecture. A comprehensive list HpaB mutants evaluated is provided in **Table S1**.

One of the theorized advantages of biosynthesizing L-DOPA over exogenous supplementation when expressing proteins containing L-DOPA is that the process itself helps maintain a cellular environment with a high ratio of L-DOPA to L-tyrosine. We hypothesized that increased activity from our *Ec*HpaB variant, GLRL-ALVH-A, would yield both a higher titer of L-DOPA than wildtype and consume more L-tyrosine, resulting in a higher ratio between the two species. To test this, we expressed a wildtype *Ec*HpaB and our GLRL-ALVH-A variant and analyzed the culture supernatant by LC-MS at two time points (**Figure 3d**). We chose an initial time point of six hours post induction of *Ec*HpaB to closely reflect the conditions of our previous fluorescent reporter assays and the 16-hour time point to represent overnight protein expression. Our results show that at six hours post induction, the GLRL-ALVH-A variant has consumed most of the L-tyrosine in the media and produced ∼6-fold more L-DOPA than wildtype. While wildtype achieved a 43:67 ratio of L-DOPA:L-tyrosine, the GLRL-ALVH-A variant reached a ratio of 66:1, representing a ∼153-fold increase. Notably, L-tyrosine levels fell below the limit of detection (<0.5 ng/mL) in both conditions by the 16-hour mark, indicating a plateau in both cellular growth and L-DOPA biosynthesis. We hypothesize that the accelerated L-tyrosine depletion observed at six hours is a key driver of greater translational incorporation of L-DOPA. We attribute the lower L-DOPA titers recorded at 16 hours to product degradation outpacing synthesis.

Variant GLRL-ALVH-A was compared to the previously reported mutants G295A and G295R, as well as to the G295C and G295T variants isolated here (**Figure S10**). While both G295A and G295R were more active than wild-type *Ec*HpaB, G295C and G295T had even higher activity, and GLRL-ALVH-A was substantially more active, consistently displaying 40-fold higher signal than wild-type in our fluorescent reporter assay. Despite significant distance between the active site cavity of HpaB and Gly 295 and L411, beneficial mutations isolated at these latter positions did not confer improvements when paired with the highly active loop mutants. In addition, we evaluated four HpaB variants previously reported to enhance the production of hydroxytyrosol for their ability to biosynthesize L-DOPA; 23F9, 23F9-M4, HpaB^TLE^ and HpaB^TLEH^ (**Figure S11**)^36–38^. None appeared to generate L-DOPA in a fluorescence assay using the MFTE-N293H variant, although HpaB^TLE^ had been experimentally determined to not accept L-tyrosine as a substrate and was not expected to be functional.

### Improved biosynthesis facilitates translational incorporation of L-DOPA

To demonstrate that an improved L-DOPA biosynthesis pathway can facilitate the downstream production of proteins containing the ncAA, we leveraged a previously developed Gen2 L-DOPA aaRS derived from the *Methanocaldococcus jannaschii* tyrosyl-tRNA synthetase. This aaRS is highly active but does not possess any editing functionality^39^. It has recently been shown to selectively bind L-DOPA and not appreciably interact with L-tyrosine *in vitro*^40^, but demonstrates only a moderate ability to discriminate between the two amino acids in living cells, and thus will charge its cognate tRNA with both L-DOPA and L-tyrosine. To quantify mischarging, a ratiometric GFP Y66DOPA S205C (designated OrGFP) fluorescent reporter can be used^41^. When L-DOPA is incorporated into the OrGFP chromophore, a unique, orange emission is observed (*λ*_ex_: 535 nm, *λ*_em_: 585 nm), while L-tyrosine incorporation yields primarily green fluorescence (*λ*_ex_: 450 nm, *λ*_em_: 500 nm) with only minor contributions from L-DOPA.

The recoded *E. coli* strain B95.ΔA was co-transformed with three plasmids encoding the orthogonal translational machinery, the OrGFP reporter, and *Ec*HpaB variants along with HpaC, respectively. Following a staged induction in which *Ec*HpaB was induced prior to expression of the OrGFP reporter to allow time for conversion of L-tyrosine to L-DOPA, fluorescence was measured (**Figure S12a**). OrGFP spectra measured from cells expressing HpaB variant GLRL-ALVH-A had the highest Orange:Green ratio and outperformed the addition of 1 mM exogenous L-DOPA to a ΔHpaB control. The variant GMRQ from the first library yielded a fluorescence ratio between wild-type *Ec*HpaB and GLRL-ALVH-A, consistent with its intermediate activity in the genetic assays.

To quantify L-DOPA incorporation, OrGFP was expressed in 200 mL cultures, purified, and analyzed by top-down mass spectrometry. The Orange:Green fluorescence ratio from the larger cultures was also measured and found to be consistent with the 1 mL assay format (**Figure S12b**). Initial analysis revealed that while the ΔHpaB control that only contained L-tyrosine had one distinct peak at the correct molecular mass, the L-DOPA-containing species had multiple peaks that were separated by 16 Da, consistent with a series of oxidation events (**Figure S12c**). To understand if this process was time dependent, we compared an OrGFP sample expressed using GLRL-ALVHA-A using measurements taken shortly after purification and after 6 months of storage at 4 °C. The 6-month-old sample showed a dramatic increase in the presence of three peaks, corresponding to successive oxidations (**Figure 4a-b**). No similar oxidation events were observed in tyrosine-only samples. To determine where this oxidation was taking place, we used UVPD to generate fragment ions spanning OrgGFP (**Fig S13**). We examined two distinct sets of fragment ions, one containing the L-tyrosine/L-DOPA incorporation site (position 65) but not the oxidized residue, and one containing only the oxidized species. While we were unable to fully resolve the identity of the oxidized species, the C-terminal fragment ions (z_49_ + 2O)^6+^ and (z_42_)^5+^ localized the oxidation to the region between His198 and Cys204. Insufficient fragment ions were recovered to confidently map the locations of the oxidized sites associated with the protein containing three oxidations.

**Figure 4:**
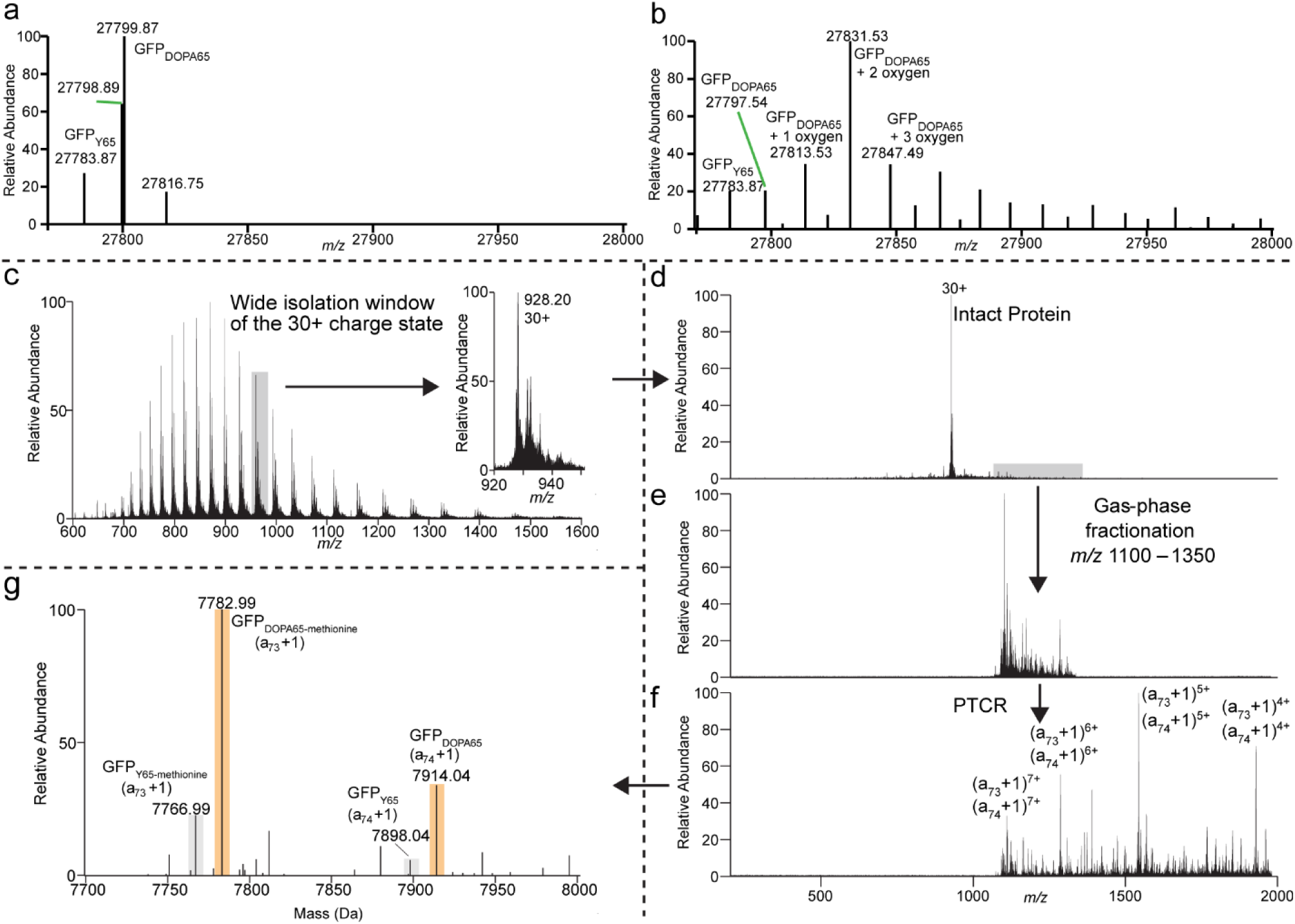
Mass spectrometry analysis of L-DOPA-containing OrGFP shows oxidation. Deconvoluted mass spectra of OrGFP_L-DOPA65_ expressed with HpaB GLRL-ALVH-A: a) before and b) after storage for 6 months at 4°C, showing oxidation of the OrGFP_DOPA65_ species. For characterization of the protein, c) ESI is used to generate the MS1 spectrum of the protein, and a selected charge state (30+, shaded in gray) is isolated, followed by d) UVPD (1 laser pulse, 0.5 mJ) to generate fragment ions, then e) isolation of all fragment ions in the window spanning *m/z* 1100-1350 (shaded in gray), followed by f) PTCR using a 6 ms reaction time. Key (a_73_ + 1) and (a_74_ + 1) ions in different charge states are labelled. g) Relative abundances are determined from the deconvoluted spectrum. The intensities of the (a_73_ + 1) ions and (a_74_ + 1) ions generated upon UVPD of the OrGFP_Y65_ variants (including both the full sequence protein and the isoform that has lost methionine) are used to estimate the portions of OrGFP_Y65_ and OrGFP_DOPA65_. Fragment peaks are shaded in gray for OrGFP_Y65_ and orange for OrGFP_DOPA65_.

To avoid confounding variables in the quantification of the protein, we used a wide ion isolation window to co-isolate all oxidation states of the protein (30+ charge state) followed by UVPD and proton transfer charge reduction (PTCR) to decongest the resulting MS/MS spectrum. The (a_73_+1) and (a_74_+1) fragment ions, which correspond to the first 73 and 74 amino acids of the protein (depending on whether the N-terminal methionine is retained), were used for quantitative analysis (**Figure 4c-g**). Using this approach, we confidently determined the percentages of L-DOPA incorporation of the purified proteins. Variant GLRL-ALVH-A, displayed 86% L-DOPA incorporation, with the remaining 14% consistent with L-tyrosine incorporation. (**Fig S12c-d**, **S14, S15**). The protein yield of OrGFP generated by this variant was 277 mg.L^-1^. The mass spectrometry results of ΔHpaB, HpaB, HpaB GMRQ, and ΔHpaB with 1 mM exogenous L-DOPA all closely matched the rank order observed in fluorescence assays with L-DOPA incorporation levels of 0%, 35%, 65%, and 73% respectively. These results are in good agreement with previously reported L-DOPA/L-tyrosine incorporation ratios using the Gen2 L-DOPA aaRS.

Given the imperfect discrimination of tyrosine and L-DOPA by the Gen2 aaRS, we hypothesized that the residual L-tyrosine incorporation was due to excess L-tyrosine present in the media and that we could achieve higher incorporation of L-DOPA if this was minimized. This is consistent with prior efforts to incorporate L-DOPA which often make use of minimal media compositions which are tyrosine-free. To test this, we repeated this experiment but used only 0.5 g.L^-1^ of yeast extract in the media (a 5-fold reduction) and observed a significant increase in L-DOPA incorporation in all conditions except in the absence of HpaB with only a minor decrease in yield **(Fig 5a-d, S16, S17)**. HpaB variant GLRL-ALVH-A achieved >97% L-DOPA incorporation and a yield of 247 mg.L^-1^. Further reductions in yeast extract were evaluated but were found to noticeably reduce cell biomass. This yield represents a significant improvement over previous efforts to biosynthesize and incorporate L-DOPA into proteins which reach similar incorporation selectivity but only yield between 3-5 mg.L^-1^ using similar reporter proteins^42,43^.

**Figure 5:**
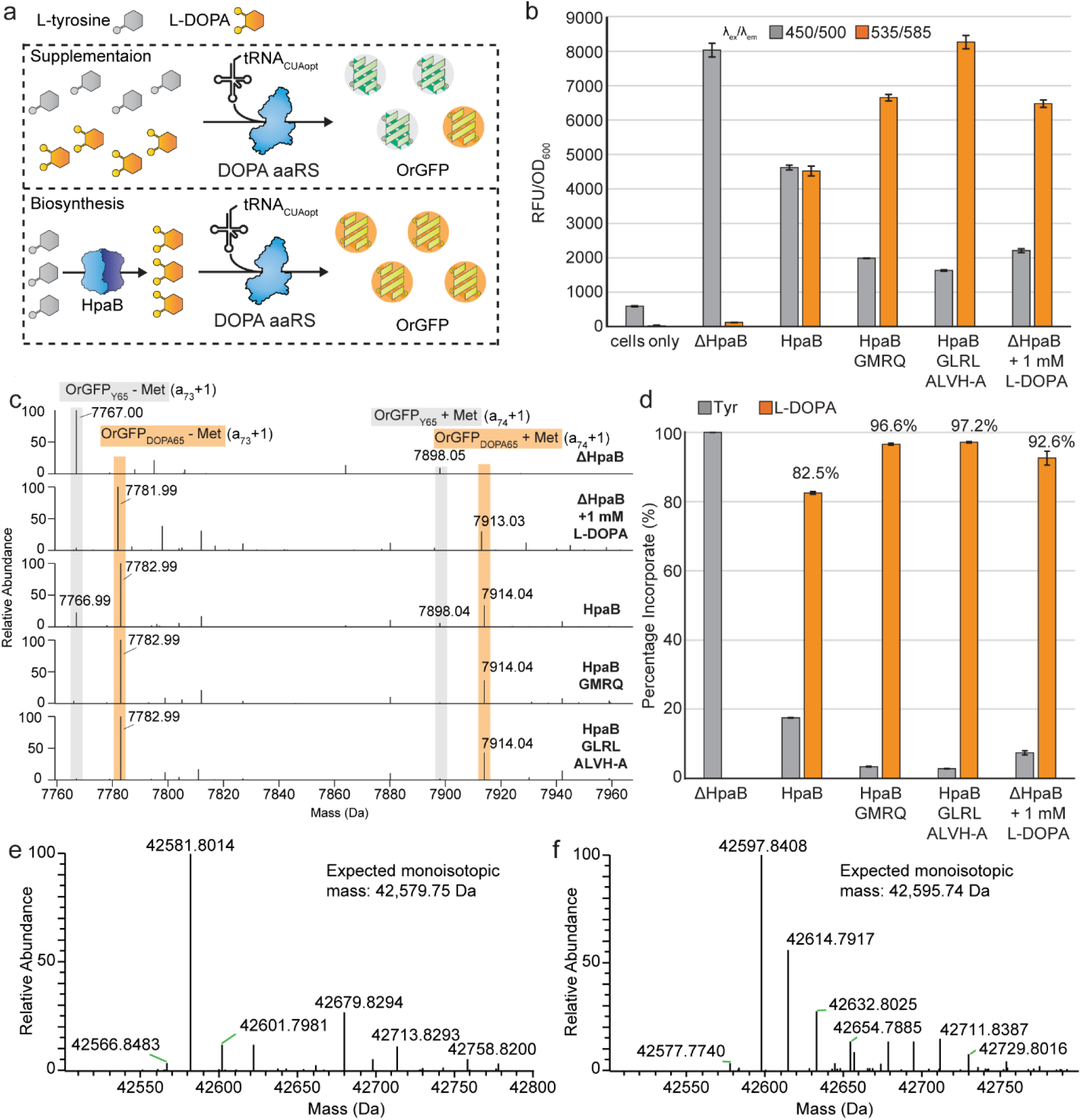
Evolved HpaB variants enhance incorporation of L-DOPA into proteins. a) Schematic of the biosynthesis and incorporation of L-DOPA into a fluorescent reporter. Co-expression of HpaB and the L-DOPA aaRS results in the translational incorporation of L-DOPA in response to a UAG codon which can be monitored by the DOPA-specific OrGFP reporter protein. b) Two channel fluorescence for the OrGFP using different variants of HpaB. ΔHpaB yields only Tyrosine incorporation and a green fluorescent signal, while the addition of 1 mM exogenous L-DOPA shifts the GFP emission towards orange. HpaB GMRQ, isolated from Library 1, represents an intermediate phenotype between our best performing HpaB variant (GLRL-ALVH-A) and the wildtype enyzme. Data represents the average fluorescence of three biological replicates normalized to cell density ± std. c) Expanded sections of deconvoluted mass spectra showing the (a_73_+1) and (a_74_+1) products generated upon UVPD (with gas-phase fractionation and PTCR) of the 30+ charge state of OrGFP for ΔHpaB, HpaB, ΔHpaB + 1 mM L-DOPA, HpaB GMRQ, and HpaB GLRL-ALVH-A. d) Relative abundances of Tyrosine- and L-DOPA-containing species were determined by mass spectrometry. The highest L-DOPA incorporation was HpaB GLRL-ALVH-A (97%) followed by HpaB GMRQ (97%), ΔHpaB + 1 mM L-DOPA (92%), HpaB (82%), and ΔHpaB (0%). Data represents the average intensity of three analytical replicates. Deconvoluted mass spectra of *Gardnerella vaginalis* ribonucleotide reductase (*Gv*RNR): e) *Gv*R2e_QSK-Tyr150_ and f) *Gv*R2e_QSK-DOPA150_.

To confirm this strategy was generalizable and could achieve a similar fidelity of DOPA incorporation outside of a model fluorescent protein, we introduced a UAG codon in place of Tyr 150 in the R2 subunit of the class Ie ribonucleotide reductase (NrdF) from the human pathogen *Gardnerella vaginalis*. Note, the taxonomic status of this species is unresolved and clusters within *Bifidobacterium*^44^. *G. vaginalis* encodes a QSK isoform of the R2e subunit which harbors a catalytic DOPA radical in the mature enzyme(*Gv*R2e_QSK_)^45^. Tyrosine oxidation to DOPA and subsequent radical formation is catalyzed by a partner enzyme NrdI which is presumed to generate an oxidizing species, possibly superoxide. In a prior report, coexpression of *nrdF* and *nrdI* in *E. coli* resulted in a heterogenous population of NrdF proteoforms containing <20% L-DOPA at residue 150. Expression of *nrdH* which encodes a small protein with thioredoxin-like activity and the R1 subunit further increased the DOPA content to ∼50%, although the respective contributions of these two proteins and how they interact with NrdI are not understood^45^. *Gv*R2e_QSK-Tyr150_ and *Gv*R2e_QSK-DOPA150_ variants were expressed using the more stringent conditions established for OrGFP and achieved yields of 123 mg.L^-1^ and 104 mg.L^-1^ respectively.

Mass spectrometry confirmed the *Gv*R2e_QSK-Tyr150_ variant contained exclusively tyrosine (**Fig 5e, S18a**), as expected for a protein expressed in the absence of NrdI. In contrast, no masses consistent with tyrosine incorporation were observed in *Gv*R2e_QSK-DOPA150_ (**Fig 5f, S18b**). A full list of the relevant deconvoluted masses for *Gv*R2e_QSK-DOPA150_ is provided in **Table S2**. This result far surpasses the level of DOPA achieved using the native site-specific post-translational machinery. An additional +17 Da species was observed for the *Gv*R2e_QSK-DOPA150_ protein which was hypothesized to represent an oxidized proteoform. However, due to the larger size of *Gv*R2e_QSK-DOPA150_ and incomplete sequence map generated by MS/MS, its location could not be determined. As shown in **(Fig S18c)**, fairly comprehensive sequence maps were generated by UVPD-MS for both engineered *RNR Gv*R2e_QSK-Tyr150_ and *Gv*R2e_QSK-DOPA150_ proteins, yielding 64% and 66% sequence coverages, respectively. These sequence coverages were obtained by combining complementary fragment ions identified from UVPD and PTCR experiments performed on precursor ions and fragment ions isolated from both lower and higher *m/z* windows, thereby maximizing sequence information across the protein backbone. Based on detection of fragment ions originating from cleavages of backbone positions that bracketed residue 150, the position of the encoded DOPA was consistent with localization at residue 150 for the *Gv*R2e_QSK-DOPA150_ protein (**Figure S18d**).

## Discussion

Transcriptional regulator-guided selection and screening is an invaluable approach for optimizing the production of L-DOPA and other metabolites. In the case of L-DOPA however, neither a transcriptional regulator nor a selection was initially available. Here, we solved this bottleneck through the discovery and engineering of a new L-DOPA-specific transcriptional regulator and biosynthetic enzyme using high-throughput emulsion-based selections. Emulsion PCR selection schemes such as CPR confer strong selective pressures and are well-suited to enriching rare variants with activity exceeding that of the parental sequence where the available dynamic range may be limiting or simply unknown^8,9^. Furthermore, the ability to easily implement a genetic inverter circuit enables exponential enrichment during counter-selection^41^. This is especially powerful for eliminating undesired variants with constitutive or promiscuous activity, a common problem during development of transcriptional regulators. Similarly, we anticipate these tools can be applied to engineer adjacent biosynthetic modules, including enzymes in the shikimate pathway needed for L-tyrosine biosynthesis, or downstream enzymes in the core BIA pathway, such as DOPA decarboxylase, using a genetic inverter. This transcriptional regulator guided approach enables large mutational landscapes to be interrogated using directed evolution and can sample complex local epistatic interactions, making it a robust platform for enzyme engineering.

We first identified a new transcriptional regulator responsive to L-DOPA, PP_2551, in *Pseudomonas putida*, optimized its minimal promoter sequence, and then engineered its ligand sensitivity using CPR. This yielded an improved PP_2551 variant designated MFTE-N293H with an EC_50_ for L-DOPA of 210 μM and 5-fold increased induction of reporter gene expression compared to the wild-type regulator. Together, these improvements represent a sensitive regulatory system capable of >100-fold induction of a fluorescent reporter when paired with native production of L-DOPA, in contrast to previously developed transcriptional regulators that report <2-fold induction at comparable L-DOPA concentrations^46^, and a fundamental improvement over traditional colorimetric outputs for screening^35,47^.

Although several different biosynthetic enzymes for L-DOPA have been shown to function in *E. coli* and other model microbial hosts for biomanufacturing^48^, extensive enzyme engineering efforts have not previously been undertaken. Leveraging the engineered L-DOPA transcriptional regulator enabled us to directly use CPR to engineer the cryptic L-DOPA biosynthetic enzyme, HpaB, a native *E. coli* hydroxylase with simple cofactor requirements. This coupling of transcriptional regulator and enzyme selection using emulsion methods had not previously been attempted, but proved to be extremely productive, likely because of the vast sequence space that could be sequentially explored. We performed iterative rounds of engineering on *E. coli* HpaB and isolated a range of substitutions that outperform previously reported HpaB mutants. Synthetic combinations of the selected variants could be rapidly screened using the transcriptional regulator for further improvements in function, and ultimately yielded the combination of GLRL-ALVH-A. This variant yielded a 6-fold higher titer of L-DOPA at six hours post-induction, which coincides with log phase window when recombinant protein synthesis is also occurring.

The large number of highly active variants that were generated by CPR also provided an opportunity to assess structure-function relationships for HpaB. We confirmed the importance of several spatially separated loci on substrate recognition, including the β32-β33 loop, the 156-159 loop, Gly 295 and Leu 411, and mapped a new beneficial mutation, T398A, in a region of HpaB not sampled in previous mutagenesis studies^35,36,49^. Clones identified from the β32-β33 loop library spanning residues G209/S210/A211/Q212 overwhelmingly preserved Gly 209 while acquiring Arg or Lys at position 211. Given the strong preference for Arg or Lys at position 211, it is unclear why the previously reported variant HpaB 23F9 containing the sequence G209/S210F/A211K/Q212F did not demonstrate activity in our fluorescence assay (**Figure S10**). This may be attributable to use of a HpaB template from a different *E. coli* strain where minor sequence variants (≥98% identity) are common.

*Ec*HpaB generally accepts an extremely broad range of substrates, and our accrued sequence substitutions can be assessed relative to previous efforts to probe substrate specificity. Mutations at Tyr 282 and Asp 284 that proved beneficial for hydroxytyrosol biosynthesis likely influence salt bridge formation between Arg 283 and Asp 219; the latter is adjacent to the β32-β33 loop and may fine-tune its flexibility, allowing the enzyme to accommodate new substrates^36,38^. Replacement of Tyr 301 and Ser 461 (within our third *Ec*HpaB library) with the corresponding residues (Phe and Ala respectively) from the *Pseudomonas aeruginosa* ortholog greatly improved hydroxylation of several polycyclic phenols^50^. Previous studies have also identified functionally important natural variation within *Ec*HpaB isoforms. Those encoding Arg 379, which directly bridges Tyr 301 and Ser 461, were capable of hydroxylating both 3-HPA and 4-HPA, whereas small residues at this position (Gly, Ser, and Cys) restricted activity to 4-HPA^51^. The small residues at position 379 also abolished activity with L-tyrosine, despite a differing placement of the hydroxyl group. Collectively, these observations inform several routes by which our CPR-based selection might be used to further alter the substrate specificity of HpaB, and wider sampling should also reveal additional cryptic loci away from the active site that alter substrate scope.

Using our engineered *Ec*HpaB (GLRL-ALVH-A) in combination with a previously developed orthogonal translation system, we were able to achieve site-specific incorporation of L-DOPA into a recombinant protein with ≥97% incorporation efficiency and an excellent overall yield of 247 mg.L^-1^. This exceeded the incorporation efficiency observed when 1 mM L-DOPA was added exogenously and represents a key step towards the production of DOPA-containing proteins at scale. Furthermore, expression of a native DOPA-containing protein, *Gv*R2e_QSK-DOPA150_, resulted in near quantitative DOPA incorporation and a yield exceeding 100 mg.L^-1^ in shake flask culture. In comparison, many prior efforts did not report yields or did not purify and characterize expressed proteins^41,52,53^. When a *Citrobacter freundii* TPL and a *MjY*-derived DOPA aaRS were used for L-DOPA incorporation only 5.09 mg.L^-1^ of protein was obtained. Similarly, when the *Mus musculus* tyrosine hydroxylase was combined with ChePheRS, a chimeric pyrrolysyl-tRNA synthetase with a phenylalanine-tRNA synthetase editing domain, this yielded only 3.1 mg.L^-1^. While in both cases there was near 100% incorporation efficiency,^42,43^ these reports utilized minimal medium devoid of L-tyrosine, which limits biomass and protein yield. To further bolster production, more investigations into defined media with precise control of L-tyrosine could yield an optimal concentration where cell viability is maintained while maximizing L-DOPA incorporation fidelity. Additionally, HpaB variants that improved the conversion of L-tyrosine to L-DOPA could be combined via a genetic fusion with *Ec*TyrB. These fusions have been shown to be functional and may facilitate substrate channeling between the two terminal steps of L-DOPA biosynthesis^54^, thereby increasing the L-DOPA:L-tyrosine ratio. Such strategies would build off of the HpaB improvements we describe here, but should also improve the production not only of DOPA-containing proteins, but of natural products, including a variety of therapeutic alkaloids^48,54,55^.

The oxidation peaks identified via mass spectrometry of the L-DOPA-incorporated OrGFP reporter represent a critical finding, necessitating a re-evaluation of both past and future analytical frameworks. While the specific site of oxidation remains to be definitively mapped, we hypothesize that Cys204 is the primary candidate due to its immediate proximity to the L-DOPA residue within the protein core. It is also the only amino acid of the ones identified known to undergo three possible oxidation events: sulfenic (-SOH), sulfinic (-SO_2_H), and sulfonic acid (-SO_3_H). Structural considerations support this model; within the β**-**barrel of OrGFP, only residues 202 and 204 are interior-facing. It is less likely that solvent-facing residues (S201 and Q203) would undergo selective oxidation only upon L-DOPA incorporation. Interestingly, the S204C mutation was originally introduced to facilitate a covalent crosslink with L-DOPA and red-shift the chromophore. While this mutation successfully induced the desired color shift, previously reported data suggests this phenotype is driven by proximity-induced non-covalent interactions rather than crosslinking. Given that OrGFP is the only known reporter protein that has a change in phenotype based on the incorporation of L-DOPA or L-tyrosine, it will likely be a valuable tool in further efforts to engineer a more selective aminoacyl-tRNA synthetase for L-DOPA. By presenting a comprehensive analysis of its mass spectrometry, we hope to aid in both increasing the interpretability of the *in vivo* measurements and enhance the reproducibility of future efforts to quantify L-DOPA incorporation using this reporter.

The modularity and simplicity of our L-DOPA biosynthetic pathway not only streamline the production of this high-value metabolite but also provide a robust complement to ongoing efforts in expanding the genetic code. The PP_2551 transcriptional regulator serves as a versatile tool for future engineering, while our large data set of HpaB mutations can inform future investigations into structural and kinetic characterization. The portability of this improved pathway allows for seamless integration into sophisticated genomic backgrounds, including highly recoded strains such as Syn61 (lacking two serine sense codons and the TAG stop codon) or rEcΔ2 (relying exclusively on TAA termination). By deploying our engineered enzymes within these specialized hosts, researchers can achieve near-quantitative L-DOPA incorporation with minimal background interference or cellular toxicity.^56,57^.

## Methods

### Molecular Cloning

All plasmids were constructed using either Gibson Assembly or a hierarchical modular cloning framework (Golden Gate). Gibson Assemblies were performed with a 1:3 ratio of backbone:insert DNA and incubated at 50 °C for 2 hours. Golden Gate cloning was performed by combining 20 fmol of each plasmid harboring a modular DNA fragment. Reactions were digested and ligated for 30 cycles (37 °C for 1 minute followed by 16 °C for 2 minutes). Assemblies were transformed into *E. coli* DH10B or DB3.1 using standard methods.

### Mutagenesis and Library Construction

Site-saturation mutagenesis of the PP_2551 transcriptional regulator of HpaB was performed by PCR using degenerate (NNS) oligonucleotide primers. DNA amplicons (1 μg) were re-circularized using Gibson Assembly (16 h) and transformed by electroporation into *E. coli* DH10B. Transformants were recovered in SOC for 1 h and a small aliquot serially diluted on a plated on solid medium to determine transformation efficiency and library coverage (minimum threshold of 10-fold coverage). The remaining culture was diluted in 200 mL of LB and cultured ON in a baffled flask followed by purification of the plasmid DNA. Supercoiled plasmid DNA encoding the different site-saturation mutagenesis libraries was subsequently transformed by electroporation into *E. coli* DH10B cells harboring the corresponding CPR plasmids. Transformation efficiency and library coverage were determined as described previously and the liquid culture containing the library was used as the input for CPR. Note, CPR plasmids for PP_2551 engineering encoded the Taq DNA polymerase, while HpaB engineering was performed using an exonuclease deficient variant of the KOD DNA polymerase.

### Compartmentalized Partnered Replication

Compartmentalized partnered replication was broadly performed as reported by Thyer and co-workers but included several changes specific to the target genes and minor updates to the method as described below^41^. Overnight cultures of *E. coli* cells transformed with PP_2551 or HpaB libraries were diluted 1:100 into 2xYT supplemented with 100 μg.mL^-1^ carbenicillin and 34 μg/mL chloramphenicol. Cultures were incubated for 2 hours to reach early-to mid-log phase at which point 1 mM L-DOPA (PP_2551) or 4 ng.mL^-1^ aTc (HpaB) was added to induce expression of the DNA polymerase. After 4 hours, 100 uL of culture was removed and cells harvested by centrifugation at 4 °C. The supernatant was removed, and the cell pellet resuspended in PCR buffer containing primers that flank the gene of interest. This resuspension was combined with oil and surfactants and mixed vigorously with a tissuelyzer at 42 hz for 4 minutes to create a water-in-oil emulsion; a detailed protocol describing emulsion preparation is provided in Abil *et al*^58^. Emulsions were aliquoted into 8-strip PCR tubes and emulsion PCR was performed.

Following emulsion-PCR, the emulsions were consolidated into 1.7 mL microcentrifuge tubes and centrifuged at 10,000 x *g* for 5 minutes to generate a discrete organic phase. The top organic layer was removed and an equal volume of chloroform/isoamyl alcohol/phenol (24:1:25) pH 8.0 added. This mixture was mixed by vortex for 2 minutes and then transferred to phase lock tubes and centrifuged at 16,000 x *g* for two minutes. Following discontinuation of the phase lock tubes by the manufacturer, 2 mL microcentrifuge tubes containing 200 μL of Dow Corning High Vacuum Grease (DC-976/ DuPont Molykote High Vacuum Grease) were adopted as a replacement. The upper aqueous layer containing the CPR amplicons was recovered and transferred to a 1.7 mL Eppendorf tube and purified using a spin-column kit (Zymo Research DCC-25). CPR amplicons were eluted in 50 μL of nuclease-free ddH_2_O. 5 μL of CPR amplicons were used as the template for a secondary PCR to generate sufficient DNA for subcloning. Amplicons from the secondary PCR were gel purified to remove any small contaminating products, and 1-5 ng of secondary used as template in a tertiary PCR reaction with nested oligonucleotide primers that introduce Gibson Assembly homology arms to enable subcloning. Products were subcloned into new expression plasmids for additional rounds of CPR or characterization using Gibson Assembly with a 1:3 molar ratio of backbone to insert, using 1 μg of plasmid backbone.

### GFP and RFP Fluorescence Assays

All GFP and RFP fluorescence assays were performed in *E. coli* strain DH10B and in M9 medium supplemented with 2.5 g.L^-1^ yeast extract (M9YE), 0.5% glycerol, and 0.5 mg.mL^-1^ of L-ascorbic acid, unless otherwise specified. Transformants were selected in biological triplicates into 96-well deep-well plates and incubated at 37 °C overnight with 900 rpm agitation on a 1.5 mm orbit. Following ON growth, cells were diluted 1:33 into fresh culture medium and incubated for two hours, after which fluorescence was induced by the addition of L-DOPA to a final concentration of 1 mM or 1.6 mM (unless specified otherwise). Following four hours of induction, cells were harvested by centrifuged at 3500 x *g* at 4 °C for 10 minutes. Supernatant was discarded and cell pellets were completely resuspended in 1 mL PBS buffer. 100 μL was transferred to a 96-well assay plate for measurement of OD_600_ and green fluorescence (475 nm *ex*, 525 nm *em*) or red fluorescence (560 nm *ex*, 610 nm *em*). For assaying PP_2551 mutants, cells were co-transformed with separate plasmids encoding PP_2551 expressed from a constitutive promoter and mScarlet-I under the control of the optimized P_PP_2552_. For assaying HpaB mutants, a reporter plasmid was constructed which expressed the evolved PP_2551 variant (MFTE N293H), mScarlet-I under the control of P_PP_2552_, and HpaC. The reporter plasmids were co-transformed with another plasmid encoding HpaB under the control of P_LtetO_. HpaB expression was induced after two hours of growth by the addition of aTc to a final concentration of 50 ng.mL^-1^.

### OrGFP Fluorescence Assays

*E coli* strain B95ΔA was co-transformed with three plasmids: the first encoding HpaB under the control of P_LtetO_ and constitutively expressed HpaC, a second encoding the Gen2 L-DOPA aaRS and tRNA under the control of P_BAD_, and the third encoding OrGFP expressed from the T7 promoter^59^. Fluorescence assays were performed as described above with differences in the induction schedule. One hour after dilution of the ON cultures, aTc was added to a final concentration of 50 ng.mL^-1^ to induce expression of HpaB, and after a further one hour, IPTG and L-arabinose were added to final concentrations of 25 μM and 0.35 % w/v respectively. For flask assays and larger scale expression of OrGFP, cells were cultured ON in 5 mL of M9YE or M9(1/5)YE in test tubes overnight before being diluted 1:200 into 200 mL of M9YE containing 1 mg.mL^-1^ L-ascorbic acid in a 500 mL baffled flask. Cells were induced as described for 96-well deep-well plate assays, except for adding aTc to a final concentration of 100 ng.mL^-1^ when OD_600_ reached 0.2, and harvesting the cells after 16 hours. A 1 mL aliquot of culture medium was centrifuged to harvest cells and perform fluorescence assays.

### Ribonucleotide Reductase Expression

Ribonucleotide Reductase was expressed using the protocol described above for OrGFP, with the exception of the OrGFP-encoding plasmid being replaced with a similar plasmid containing the *Bv*R2e_QSK-TAG150_ under the control of a T7 promoter.

### Protein Purification

Cells expressing OrGFP or *Bv*R2e_QSK-TAG150_ were harvested by centrifugation and resuspended in IMAC Resuspension Buffer (50 mM NaH_2_PO_4_, 100 mM NaCl, 1 mM EDTA, pH 8.0) and lysozyme was added at a concentration of 3 mg.mL^-1^. The cells were incubated at 37 °C for 60 minutes and then lysed by sonication. Lysate was clarified by centrifugation and the EDTA quenched by addition of MgCl_2_ at a final concentration of 10 mM. The supernatant was purified using Ni-NTA resin and eluted with IMAC elution buffer (50 mM NaH_2_PO_4_, 300 mM NaCl, 300 mM imidazole, pH 8.0). Proteins were dialyzed into a sodium phosphate buffer (20 mM NaH_2_PO_4,_ 50 mM NaCl, and pH 7.4) for storage and further characterization.

### HPLC-ECD

Plasmids encoding WT HpaB, evolved HpaB variants, or inactive controls were transformed into *E. coli* DH10B. Colonies were inoculated in biological quadruplicates into 1 mL of 2xYT supplemented with 100 μg.mL^-1^ carbenicillin and incubated for 16 h at 37 °C in a 96-well deep-well plate. The next day, cultures were diluted 1:100 into 1 mL fresh media supplemented with 10 mM L-ascorbic acid and cultured for six hours. 500 μL of culture media was clarified by centrifugation at 3000 x *g* for seven minutes. The supernatant was then transferred to a 0.2 μm filter tube (EMD Millipore: UFC30GVNB) and centrifuged at 12,000 x *g* for three minutes. 50 μL of filtered supernatant was diluted with 450 μL of MD-TM mobile phase (Thermo: 70-1332) and 10 μL aliquots used for HPLC-ECD measurements. The HPLC-ECD measurements were performed on a Thermo UltiMate 3000 system with an autosampler (Thermo: WPS-3000TBRS) paired to a Dionex UltiMate 3000 electrochemical detector (Thermo: ECD-3000RS) using a coulometric ECD cell (Thermo: 6011-RS). Sampling injections were passed through a 2.1 x 50 mm, 5 µm C18 reverse phase column (Agilent Eclipse plus: 9597 46-902) incubated at 30 °C with a flow rate of 0.5 mL/min. The applied potential of the ECD cell was 300 mV vs. Pd with a 10 Hz collection rate. Fresh L-DOPA standards from 12.5 to 100 pmol were prepared and measured during 7-minute runs with standard peaks observed at ∼2.1 minutes. Data was visualized and analyzed with Chromeleon 7 CDS software.

### Mass Spectrometry

OrGFP and *Gv*R2e_QSK_ proteins were diluted to 10 μM and desalted using P6 Micro Bio-Spin columns for OrgGFP and 10 kDa MWCO Amicon Ultra centrifugal filters for *Gv*R2e_QSK_ into 50/50 methanol/water with 0.1% formic acid. Proteins solutions were analyzed on a Thermo Orbitrap Eclipse mass spectrometer (Thermo Fisher, San Jose, CA) modified for UVPD^60^. Samples were introduced by nanoelectrospray ionization using Au/Pd-coated borosilicate static tips (OD 1.2 mm, ID 0.69 mm) pulled in-house. A voltage of 1.4 kV was applied for ESI. All spectra were collected at 240K resolution using 200 averages in full profile mode and intact protein mode. Spectra were deconvoluted with Xtract in Freestyle using a S/N of 10.

For the MS^2^ analysis, the 30+ charge state of OrGFP or the 43+ charge state of *Gv*R2e_QSK_ was isolated and characterized by 193 nm UVPD using 1 laser pulse at 0.5 mJ per pulse. Gas-phase fractionation was performed during a two window pseudo-MS^3^ procedure in which all fragment ions generated by UVPD that were lower in *m/z* than the precursor were isolated and subjected to proton transfer charge reduction and then all fragment ions produced by UVPD that were higher in *m/z* than the precursor were subjected to proton transfer charge reduction. For OrGFP, the lower *m/z* window spanned *m/z* 200 – 880 and the higher *m/z* window spanned *m/z* 960 – 2000, whereas for *Gv*R2e_QSK_, the lower *m/z* window spanned *m/z* 110–930 and the higher m/z window spanned *m/z* 1015– 1985. In addition, a narrower window spanning only *m/z* 1100-1350 for OrGFP was used to capture multiple charge states of the (a_73_+1) ion. Proton transfer charge reduction (PTCR) was performed using a 6 ms reaction time with perfluoroperhydrophenanthrene (PFPP) used to generate the proton-scavenging reagent anions. All spectra were collected in triplicate. For OrGFP, the oxidation site was localized using fragment ions containing a modification of +15.99 Da, corresponding to one oxidation. Semi-quantitative analysis of L-DOPA incorporation in OrGFP was done using the relative abundances of the (a_73_+1) ion with the loss of the N-terminal methionine and (a_74_ +1) for GFP without the loss of the N-terminal methionine.

### LC-MS Analysis of L-Tyrosine and L-DOPA

Samples for LC-MS were prepared using *E. coli* strain DH10B containing plasmids expressing either wild-type *Ec*HpaB or the best engineered variant (GLRL-ALVH-A) under the inducible control of P_LtetO_ with HpaC constitutively expressed. Cells containing an empty plasmid were used for a ΔHpaB control. Overnight cultures were diluted 1:200 into 200 mL of M9 minimal media supplemented with 2.5 g.L^-1^ yeast extract, 0.5% glycerol, and 1 mg.mL^-1^ ascorbic acid. The cells were agitated at 37 °C until an OD_600_ of 0.2 was reached followed by addition of aTc to a final concentration of 100 ng.mL^-1^. Aliquots of 5 mL were taken after six and 16 hours post induction and clarified via centrifugation prior to storage at 4 °C until LC-MS analysis.

LC-MS analysis of tyrosine and L-DOPA was performed on an Agilent 6470 Triple Quadrupole LC/MS System interfaced to an Agilent 1290 Infinity II UPLC System through an Agilent Jet Stream electrospray ionization source operated in positive ionization mode. The LC mobile phases consisted of A (0.2% formic acid in water) and B (acetonitrile). Separations were carried out using a Waters Acquity Premier HSS T3 column (2.1 mm ID × 100 mm, 1.8 µm particle size) operated at a flow rate of 0.4 mL/min and a column temperature of 25 °C. The LC gradient was as follows: 0–4.5 min, 100% A; ramped to 90% B at 6.5 min; held at 90% B until 9 min; followed by re-equilibration to initial conditions for 3 min prior to the next injection. The sample injection volume was 25 µL. ESI source parameters were optimized as follows: gas temperature 350 °C, gas flow 6 L/min, nebulizer pressure 35 psi, sheath gas temperature 400 °C, sheath gas flow 12 L/min, capillary voltage 3000 V, and nozzle voltage 500 V. MRM parameters for L-DOPA, tyrosine, and the ^13^C-labeled tyrosine internal standard are reported in Table 1.

**Table 1:** MRM parameters for L-DOPA, L-Tyrosine, and ^13^C-Tyrosine.

| Cpd Name | Precursor Ion | Product Ion | Dwell Time (ms) | Frag (V) | CE (V) | Cell Acc (V) |
| --- | --- | --- | --- | --- | --- | --- |
| L-DOPA | 198.1 | 107.1 | 50 | 94 | 29 | 4 |
| L-DOPA | 198.1 | 77.1 | 50 | 94 | 53 | 4 |
| Tyrosine | 182.1 | 91.1 | 50 | 90 | 33 | 4 |
| Tyrosine | 182.1 | 77.1 | 50 | 90 | 49 | 4 |
| Tyrosine-<sup>13</sup>C-IS | 192.1 | 98.1 | 50 | 90 | 49 | 4 |
| Tyrosine-<sup>13</sup>C-IS | 192.1 | 83 | 50 | 90 | 49 | 4 |

Quantitative data analysis was performed using Agilent MassHunter Quantitative Analysis. The lower limit of quantitation (LLOQ) for both tyrosine and L-DOPA was 0.5 ng/mL, with a linear dynamic range extending to 100 ng/mL. The *R*² values for L-DOPA and tyrosine were 0.9998 and 0.9996, respectively. Quality control recoveries ranged from 98.5% to 100.5% for L-DOPA and from 95.4% to 99.4% for tyrosine.

### Replicates and Statistics

All experiments were performed in biological and technical triplicate unless specifically indicated. Fluorescence data is plotted as the mean ± std, unless specifically stated otherwise.

## Data Availability

Relevant DNA sequences can be found in Supplementary Tables S3 and S4. Materials, reagents, and detailed protocols are available from the authors upon request. All top-down mass spectrometry data has been deposited into the MassIVE public repository with the identifier MSV000103442.

## Acknowledgements

Funding from the NIH (5K99CA207870-02 to R.T., R01EB026533 to A.D.E, R35GM139658 to J.S.B.), the Welch Foundation (C-2167-20230405 to R.T., F1155 to J.S.B.), and Ajinomoto Co., Inc. (UTA16-001150) are acknowledged. The authors also acknowledge the help of Nancy A. Moran.

## Author Information

### Authors and Affiliations

**Systems, Synthetic, and Physical Biology, Rice University, Houston, TX**

Andrew R. Gilmour

**Department of Bioengineering, Rice University, Houston, TX**

Qiyao Wei

**Department of Molecular Biosciences, The University of Texas at Austin, Austin, TX**

Chad Wang, Vidya Pandarinath, Simon d’Oelsnitz, and Andrew D. Ellington

**Department of Chemistry, The University of Texas at Austin, Austin, TX**

Jessica Hellinger, Prithvijeet Singh Kharnal and Jennifer S. Brodbelt

**Department of Integrative Biology, The University of Texas at Austin, Austin, TX**

Eli Powell

**Department of Chemical and Biomolecular Engineering, Rice University, Houston, TX**

Ross Thyer

### Contributions

A.R.G., Q.W., V.P. and R.T. characterized and engineered the transcriptional regulator. A.R.G., Q.W., C.W., S.D., and R.T. engineered and characterized the HpaB enzyme variants. A.R.G and Q.W. expressed and purified the proteins. E.P. and N.A.M. performed the ECD-HPLC characterization. J.H. and J.S.B performed the mass spectrometry experiments. A.R.G., A.D.E, and R.T. wrote the manuscript with input from the authors.

### Corresponding author

Correspondence to Ross Thyer.

### Ethics Declarations

A.R.G., C.W., A.D.E, and R.T. are coinventors on US provisional patent related to L-DOPA biosynthesis. The other authors declare no competing interests.

