## Supplementary Information for "Directed evolution of L-DOPA sensing and production enables efficient incorporation of an expanded genetic code into proteins"

This file contains:

18 Supplementary Figures

4 Supplementary Tables

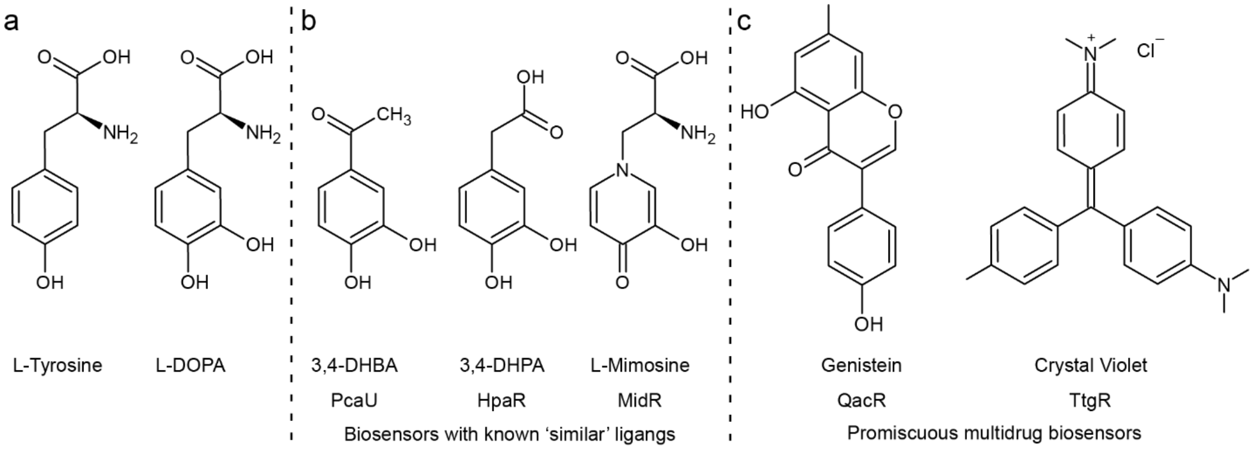


**Figure S1.** Ligand structures of known biosensors that were screened for activity on L-DOPA. a) Chemical structures of L-tyrosine and L-DOPA for reference against other ligands. b) Chemical structures of the ligands for PcaU, HpaR, and MidR. These ligands share a catechol or catechol-like structure which motivated our interest in testing their respective biosensors activity on L-DOPA. c) Chemical structures of the ligands for QacR and TtgR. These biosensors have been characterized as highly promiscuous and were tested to see if they had basal activity for L-DOPA.

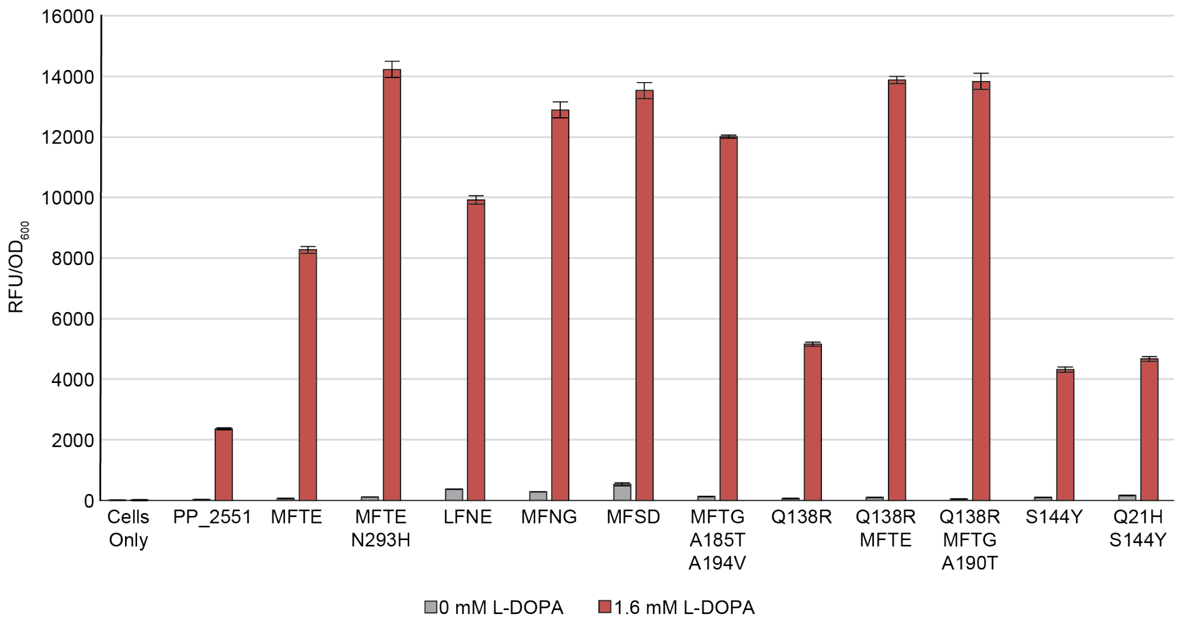


**Figure S2.** Screening of PP2551 mutations isolated following three rounds of CPR selection. PP2551 mutants were assayed using a mScarlet-I reporter under the control of the *PP2552* promoter ± 1.6 mM L-DOPA. Data represents the average fluorescence (Relative Fluorescence Units or RFU) of three biological replicates normalized to cell density, as measured by absorbance at $\lambda$ = 600 nm, ± std.

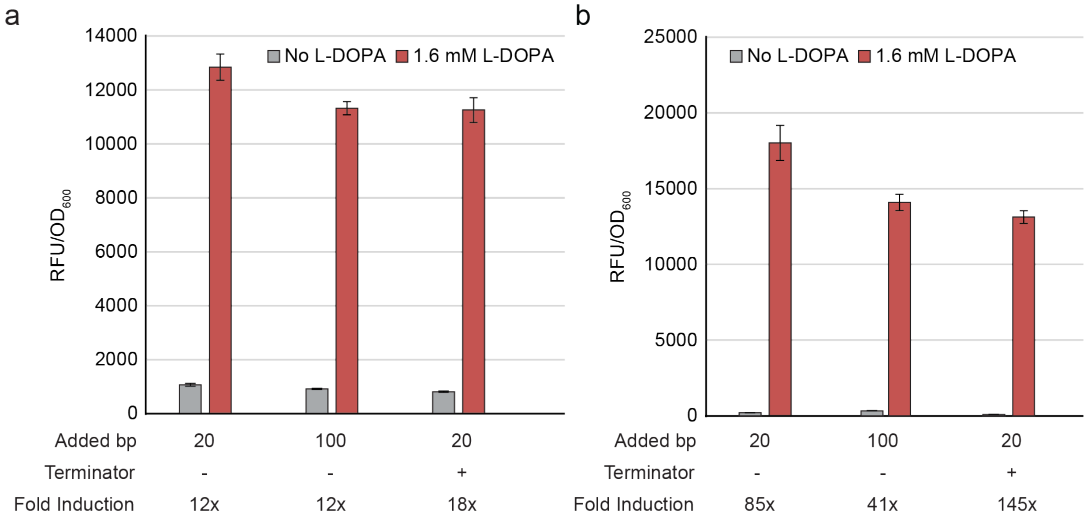


**Figure S3:** Comparison of different culture media on PP2551 biosensor promoter activity. (a) Screening of insulated promoter variants , either containing 20 or 100 base pairs (bp) of added upstream sequence and/or a terminator, using an mScarlet-I reporter and the engineered PP2551 biosensor in 2XYT media. (b) Screening of the same insulated promoter variants and biosensor but in M9 media supplemented with 2.5 g/L yeast extract. Data represents the

a

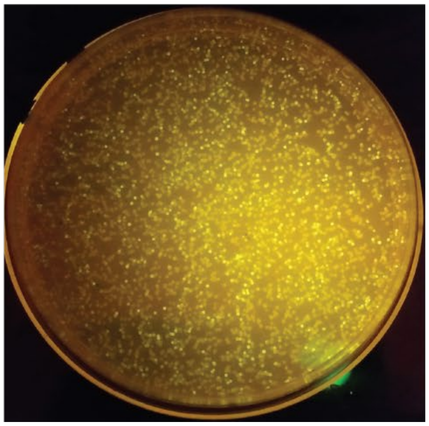

b

| 295 | 411 | Count |
|---|---|---|
| G | L | 0 |
| V | L | 3 |
| V | I | 2 |
| V | V | 2 |
| C | L | 3 |
| C | V | 3 |
| C | F | 1 |
| T | L | 2 |
| T | I | 2 |
| S | L | 1 |
| S | V | 1 |
| M | L | 2 |

c

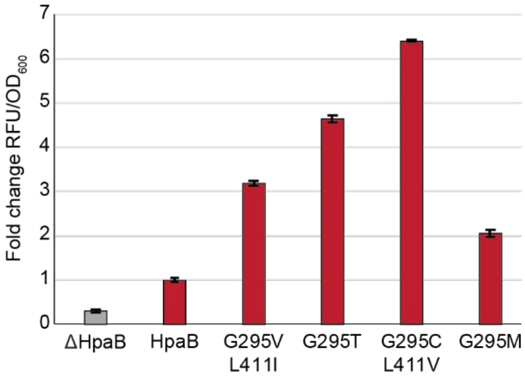


**Figure S4:** Screening of HpaB site saturation library at residues G295-L411. (a) Representative agar plate containing *E. coli* cells that have been co-transformed with a site saturation library of HpaB at G295 and L411 and a reporter plasmid with the MFTE-N293H variant of PP2551 and sfGFP. (b) Table showing the mutations found at residues G295 and L411 from 24 highly fluorescent colonies. Note, two of 24 clones failed to sequence correctly. (c) Fluorescence assay of selected variants isolated from the library. Data represents the average fluorescence of three biological replicates normalized to cell density ± std.

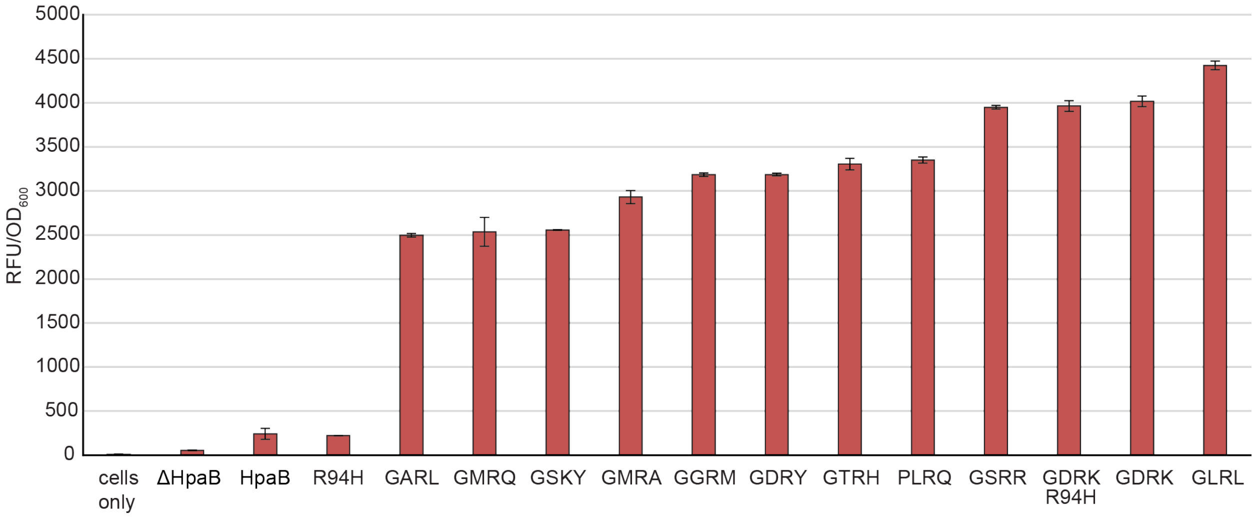


**Figure S5:** Fluorescent assay of HpaB variants isolated following three rounds of CPR. Unless otherwise noted, all mutations span residues G209/S210/A211/Q212. Mutations isolated from the first library selection via CPR were subcloned and evaluated using a reporter plasmid encoding PP2551-MFTE-N293H and mScarlet-I. Data represents the average fluorescence of three biological replicates normalized to cell density ± std. A complete list of the mutations in each variant is provided in SI Table 1.

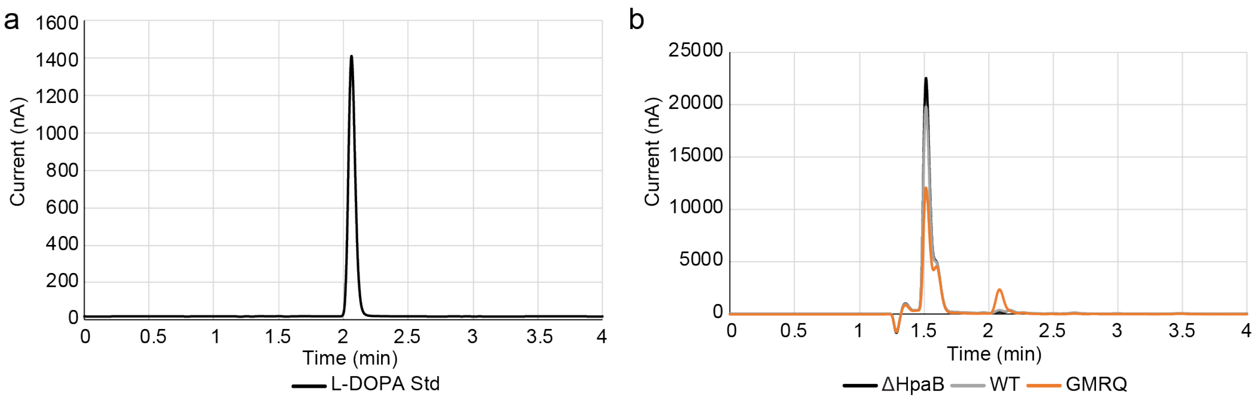


**Figure S6:** Validation of L-DOPA production from HpaB library variants using High-Performance Liquid Chromatography with Electrochemical Detection (HPLC-ECD). (a) HPLC trace of L-DOPA standard. (b) HPLC trace of culture medium from *E. coli* expressing containing a truncated HpaB (ΔHpaB), wildtype HpaB, or the HpaB-GMRQ mutant isolated from the initial CPR library selection.

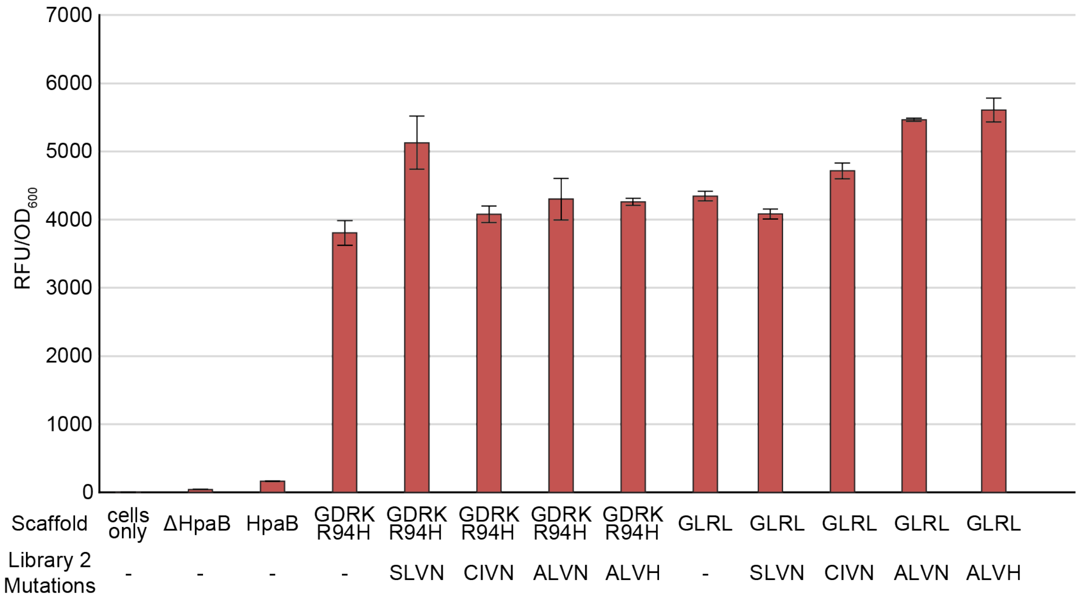


**Figure S7:** Fluorescent assays of HpaB variants isolated from the second site-saturation library. Fluorescence assay of the two best performing variants from library one (R94H/G209/S210D/A211R/Q212K and G209/S210L/A211R/Q212L) with the additional mutations isolated from library two (A156/I157/V158/N159). Variants were evaluated using a reporter plasmid encoding PP2551-MFTE-N293H and mScarlet-I. Data represents the average fluorescence of three biological replicates normalized to cell density ± std.

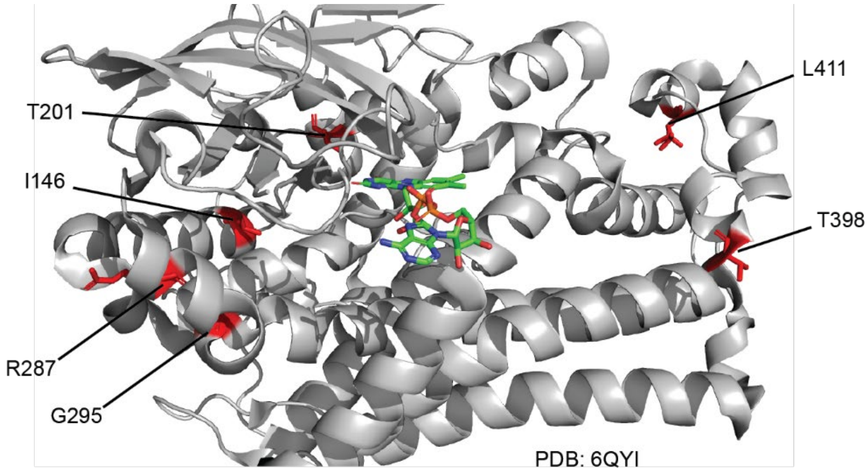


**Figure S8:** Crystal structure of HpaB showing the location of off-target and previously reported mutations in relation to the active site cavity (marked in red) and the FAD cofactor (green). Residues G295 and L411 were reported previously by Fordjour *et al.* and residues I146/T201/R287/T398 were identified in this work[1].

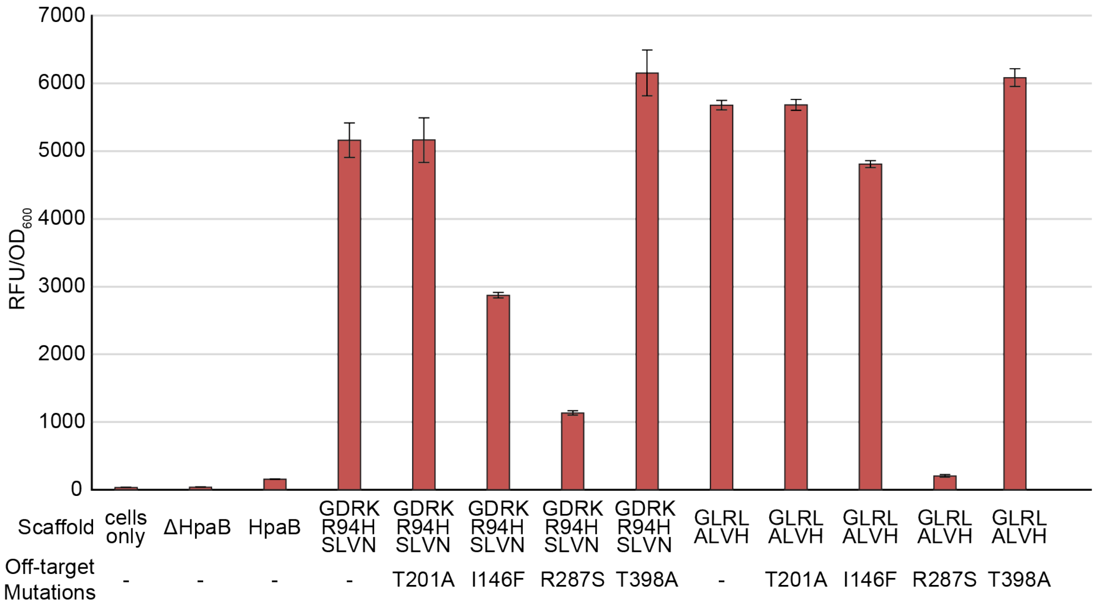


**Figure S9:** Fluorescent assays of HpaB variants with four off-target mutations. Fluorescence assay of the two best performing variants from the first library selection, GDRK-SLVN-R94H and GLRL-ALVH, with four additional off-target mutations recovered from libraries one and two. Variants were evaluated using a reporter plasmid encoding PP2551-MFTE-N293H and mScarlet-I. Data represents the average fluorescence of three biological replicates normalized to cell density ± std.

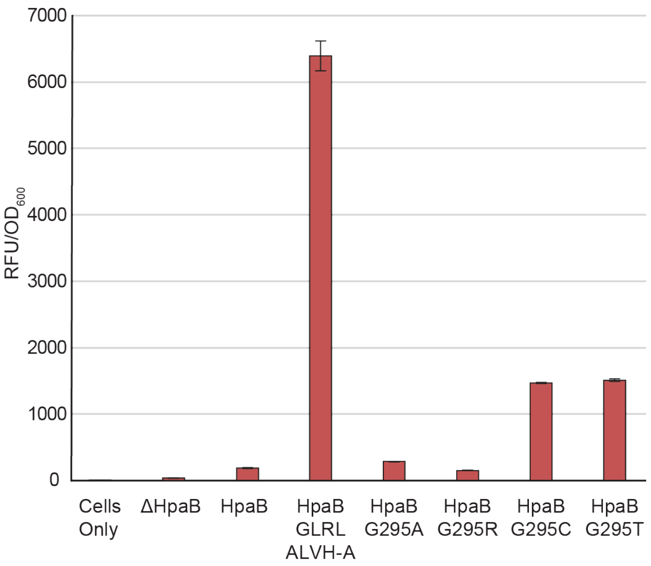


**Figure S10:** Fluorescence assay of the highest performing HpaB mutant (GLRL-ALVH-A) against mutations reported at G295 in literature (G295A and G295R) and from this work (G295C and G295T). Data represents the average fluorescence of three biological replicates normalized to cell density ± std.

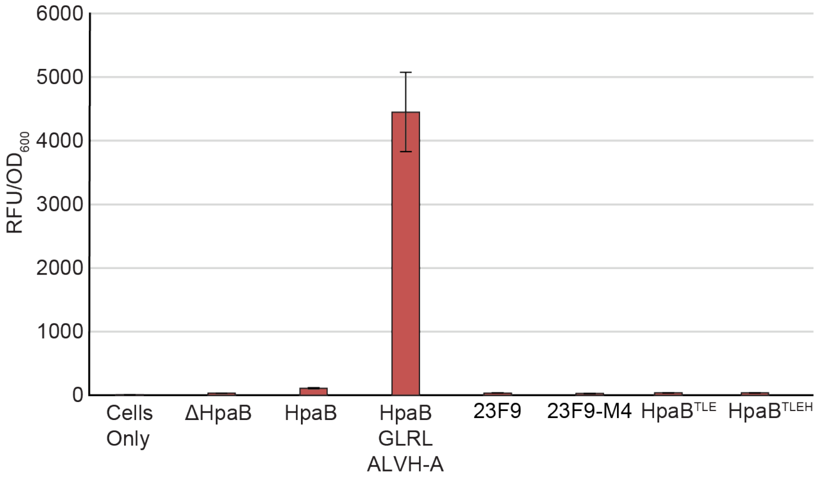


**Figure S11:** Mutations that increase HpaB activity for hydroxytyrosol production do not improve L-DOPA production. Four HpaB variants were assayed with a reporter plasmid encoding the PP2551-MFTE-N293H biosensor and mScarlet-I. 23F9 and 23F9-M4 contain mutations S210F/A211K/Q212F with 23F9-M4 containing the additional mutations T15P and D284E. HpaB$^{TLE}$ and HpaB$^{TLEH}$ both contain the mutations S210T/A211L/Q212E with HpaB$^{TLEH}$ containing the additional mutation Y282H. Data represents the average fluorescence of three biological replicates normalized to cell density ± std.

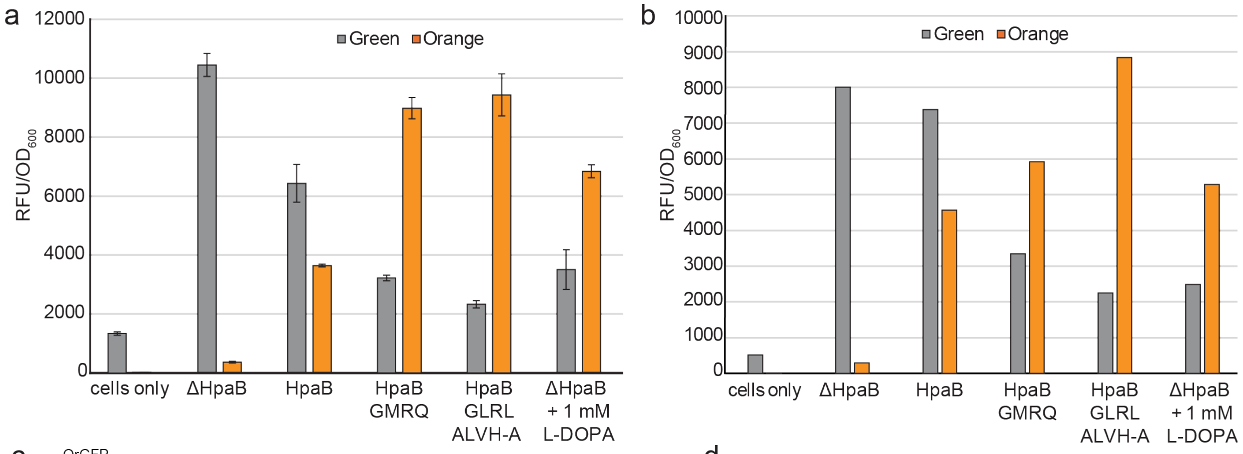


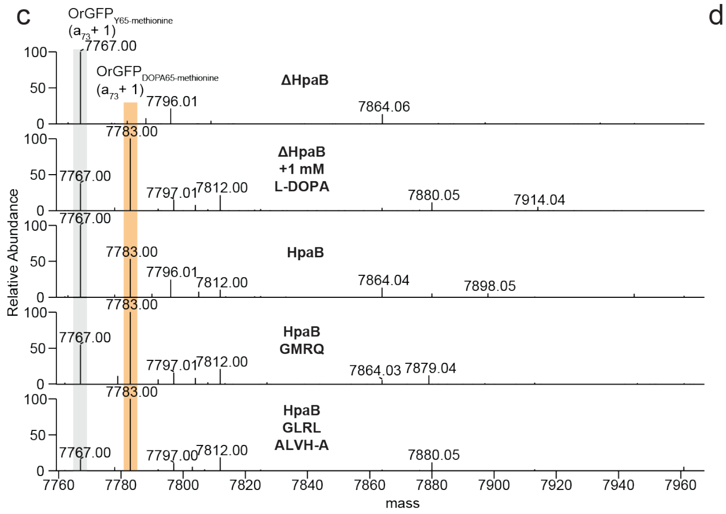


d

| Condition | % L-tyrosine | % L-DOPA |
|---|---|---|
| ΔHpaB | 100% | |
| HpaB | 65.3% ± 0.4% | 34.7% ± 0.1% |
| HpaB GMRQ | 35.5% ± 0.1% | 64.5% ± 0.3% |
| HpaB GLRL ALVH-A | 13.6% ± 0.1% | 86.4% ± 0.1% |
| ΔHpaB +1 mM L-DOPA | 27.4% ± 0.1% | 72.6% ± 0.2% |

**Figure S12:** a) Two channel fluorescence for the OrGFP using different variants of HpaB. ΔHpaB yields only tyrosine incorporation and a green fluorescent signal, while the addition of 1 mM exogenous L-DOPA shifts the GFP emission towards orange. HpaB GMRQ, isolated from Library 1, represents an intermediate phenotype between our best performing HpaB variant (GLRL-ALVH-A) and the wildtype enzyme. Green fluorescence was measured with an $\lambda_{ex}$ of 450 nm and an $\lambda_{em}$ of 500 nm, and red fluorescence was measured with an $\lambda_{ex}$ of: 535 nm and an $\lambda_{em}$ of 585 nm. Data represents the average fluorescence of three biological replicates normalized to cell density ± std. b) Fluorescence from baffled flasks expressing Orange-GFP with different HpaB variants prior to purification. c) Deconvoluted mass spectra showing the ($a_{73}$+1) products generated upon UVPD (with gas-phase fractionation and PTCR) of the 30+ charge state of GFP. Fragment peaks are shaded in gray for $OrGFP_{Y65}$ and orange for $OrGFP_{L\text{-}DOPA65}$, each observed with the loss of the N-terminal methionine. d) The relative percentages of L-DOPA incorporation.

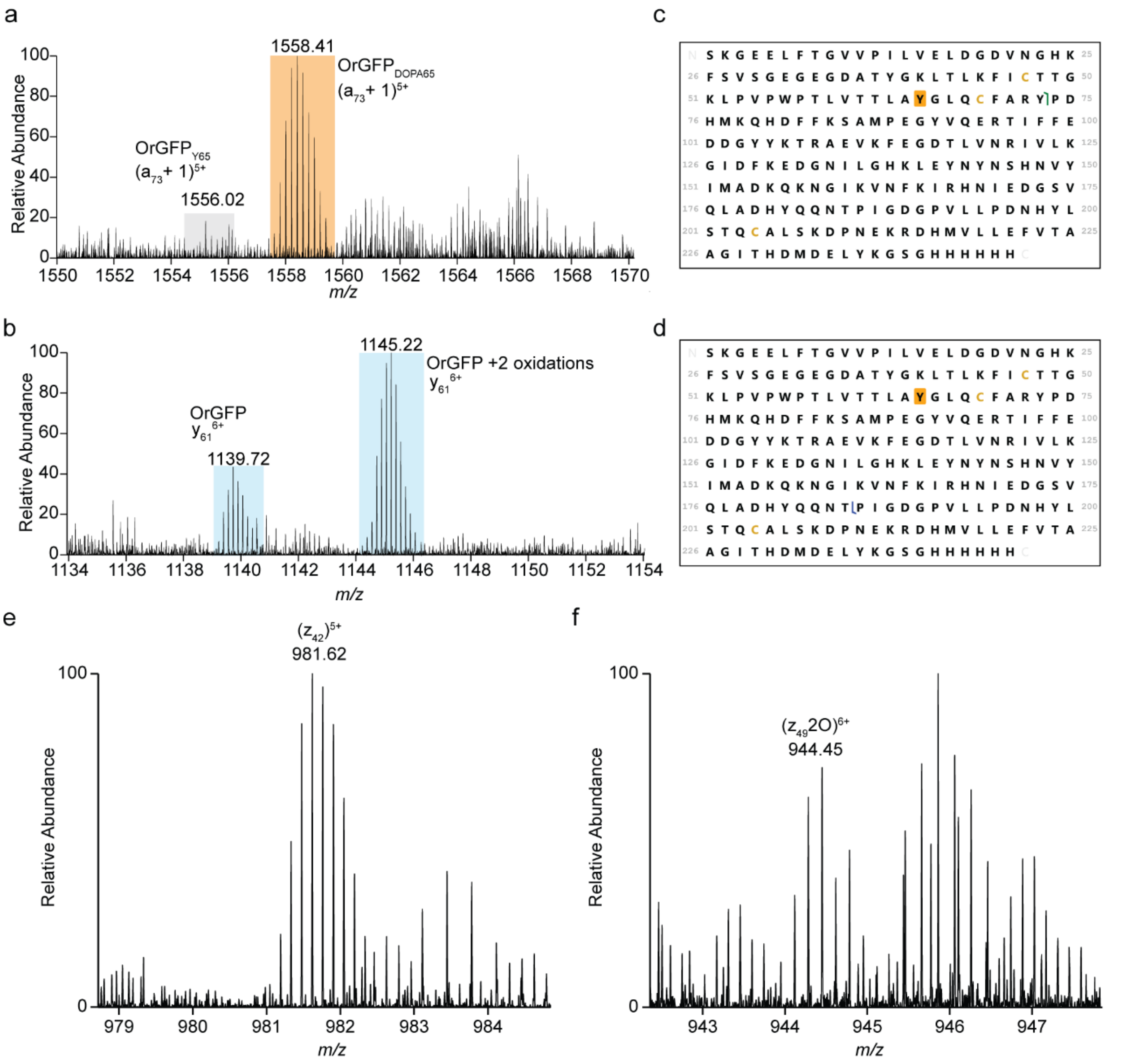


**Figure S13:** Sections of UVPD mass spectra of the 30+ charge state of OrGFP for the GLRL-ALVH-A sample after storage for six months at 4 °C showing: a) $(a_{73}+1)^{5+}$ ions for the Tyr65 and L-DOPA proteoforms, and b) $y_{61}^{6+}$ ions with proteoforms containing 0 and 2 oxidations. Fragment peaks are shaded in gray for $OrGFP_{Y65}$, orange for $OrGFP_{DOPA65}$, and blue for $OrGFP_{DOPA65}$ oxidized proteoforms, c) Sequence maps showing the backbone cleavage sites from which the site-localizing $a_{73}$ ions (cleavage between Y73 and P74) and d) $y_{61}$ ions (cleavage between Tyr185 and Pro186) originate. Tyr65 is shaded in gold to indicate that it represents the position where DOPA65 replaces Tyr65 for the engineered proteoform. Expanded sections of the UVPD mass spectrum of the 30+ charge state of OrGFP for the GLRL-ALVH-A sample after storage for six months at 4 °C showing: e) standard $z_{42}^{5+}$ ion and f) $(z_{49}+2O)^{6+}$ localizing the oxidations between His198 and Cys204.

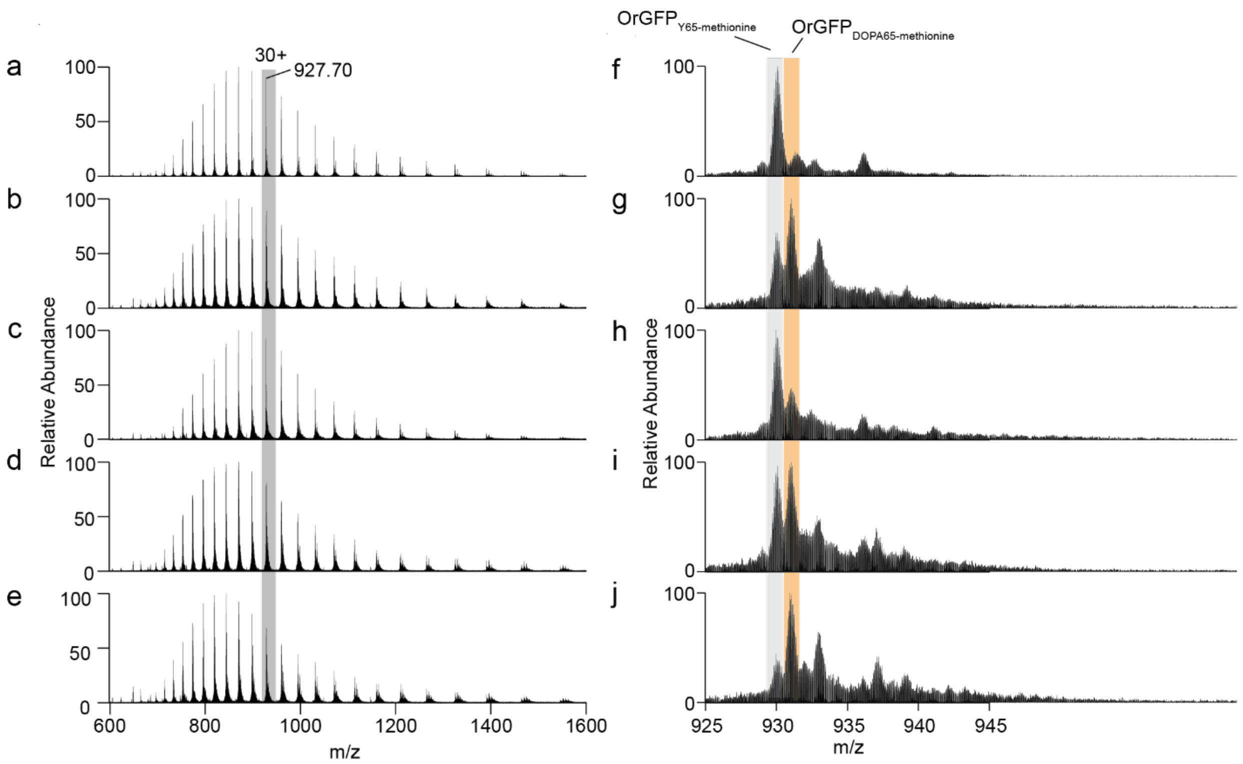


**Figure S14:** MS1 spectra of OrGFP proteins from five samples: a) ΔHpaB, b) ΔHpaB + 1 mM L-DOPA, c) HpaB, d) HpaB GMRQ and e) GLRL-ALVH-A with f-j) expanded sections of the 30+ charge state of OrGFP for each sample. Ion peaks are shaded in gray for $OrGFP_{Y65}$ and orange for $OrGFP_{DOPA65}$, each observed with the loss of the N-terminal methionine.

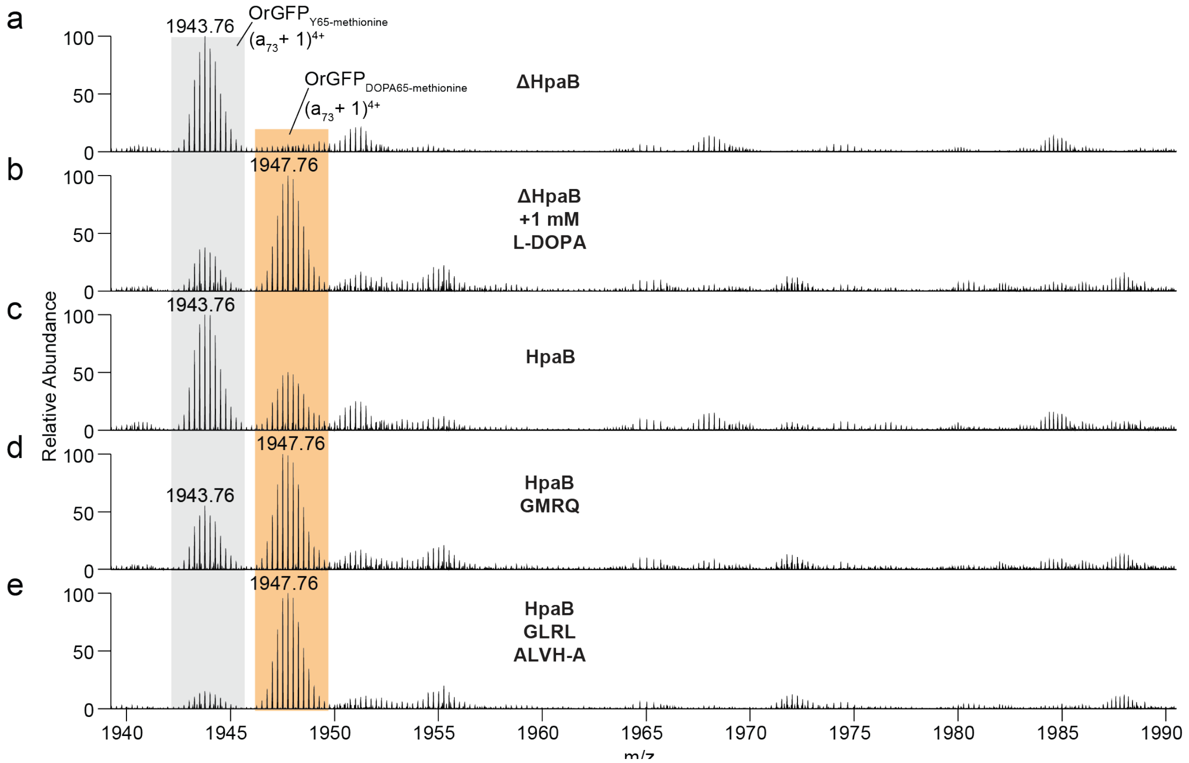


**Figure S15:** Expanded sections of mass spectra showing the $(a_{73}+1)^{4+}$ product ions generated upon UVPD (with gas-phase fractionation and PTCR) of the 30+ charge state of GFP of the following samples: a) ΔHpaB, b) ΔHpaB + 1 mM L-DOPA, c) HpaB, d) HpaB GMRQ and e) GLRL-ALVH-A. Ion peaks are shaded in gray for OrGFP$_{Y65}$ and orange for OrGFP$_{L\text{-}DOPA65}$, each observed with the loss of the N-terminal methionine. The relative abundances of the two highlighted fragment ions are used for quantification of L-DOPA incorporation.

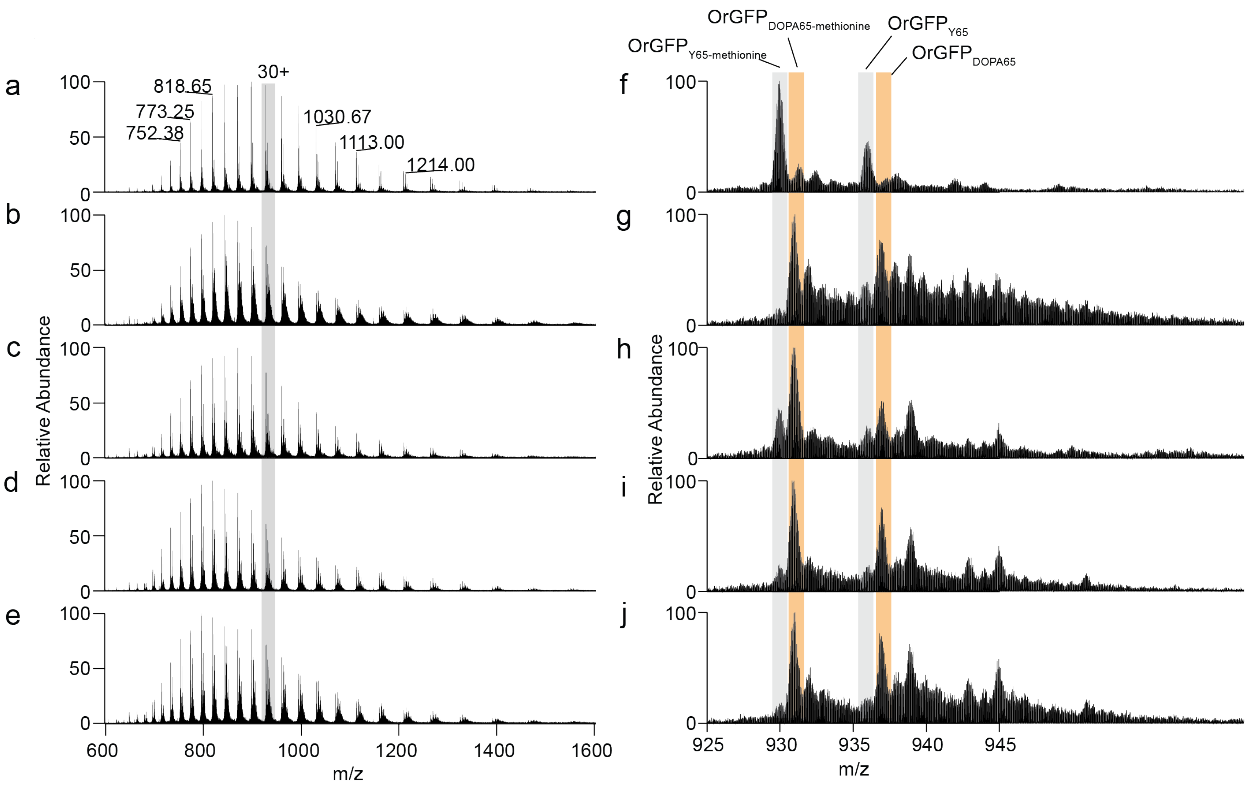


**Figure S16:** MS1 spectra of OrGFP proteins from 5 samples: a) ΔHpaB, b) ΔHpaB + 1 mM L-DOPA, c) HpaB, d) HpaB GMRQ and e) GLRL-ALVH-A with f-j) expanded sections of the 30+ charge state of the OrGFP for each sample. Ion peaks are shaded in gray for $OrGFP_{Y65}$ and orange for $OrGFP_{L\text{-}DOPA65}$, observed with and without the loss of the N-terminal methionine.

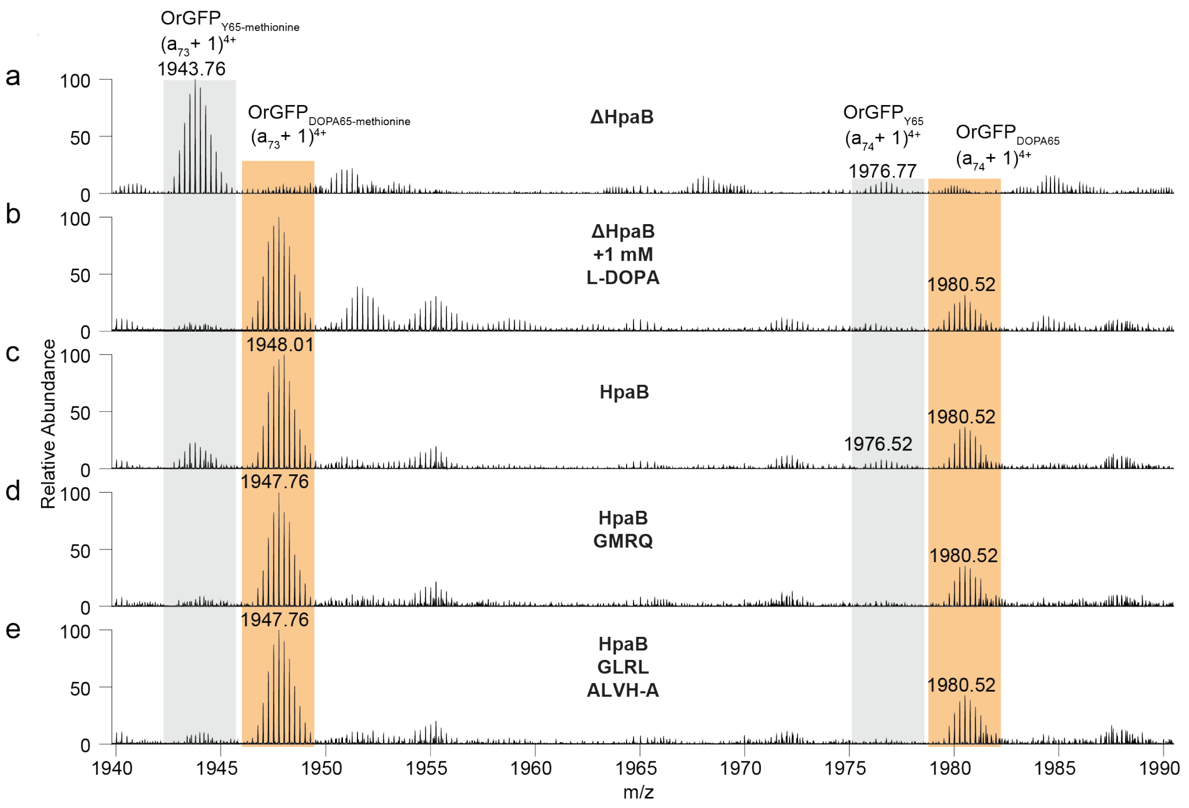


**Figure S17:** Expanded sections of deconvoluted mass spectra showing the ($a_{73}$+1) and ($a_{74}$+1) products generated upon UVPD (with gas-phase fractionation and PTCR) of the 30+ charge state of OrGFP from samples: a) ΔHpaB, b) ΔHpaB + 1 mM L-DOPA, c) HpaB, d) HpaB GMRQ and e) GLRL-ALVH-A. Ion peaks are shaded in gray for $OrGFP_{Y65}$ and orange for $OrGFP_{L\text{-}DOPA65}$, each observed with the loss of the N-terminal methionine. The relative abundances of the two highlighted fragment ions are used for quantification of L-DOPA incorporation.

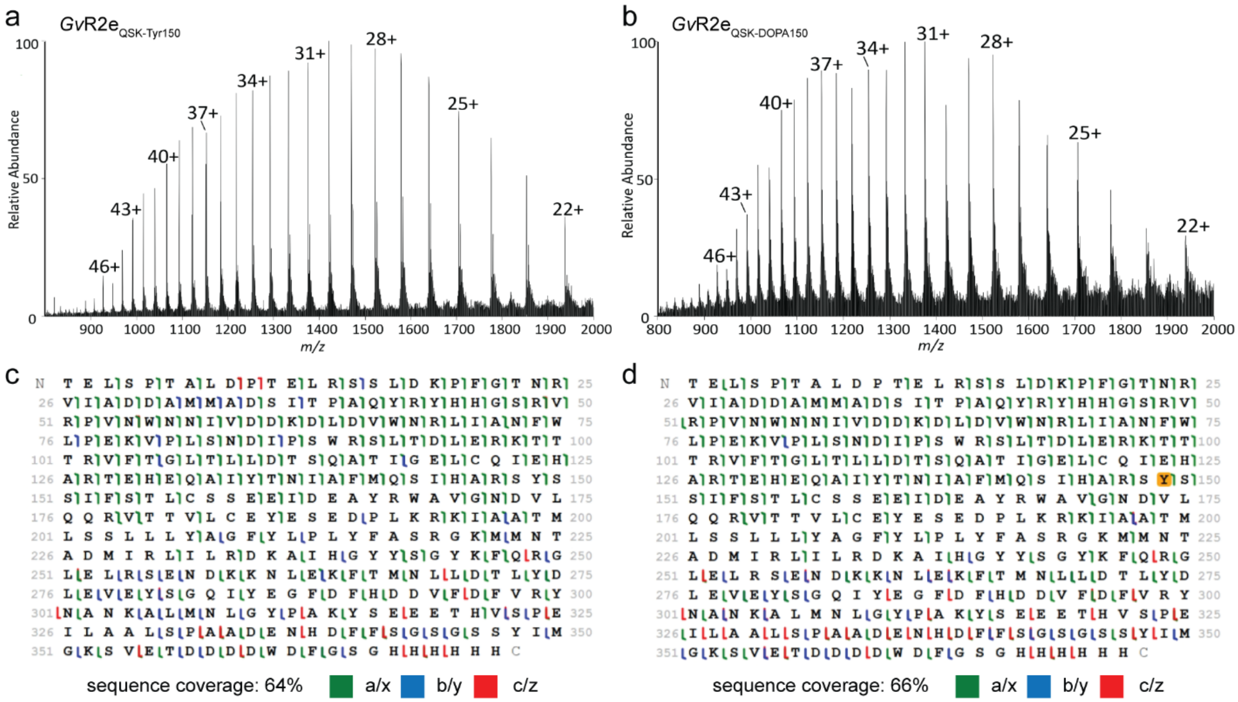


**Figure S18:** MS1 spectra of *Gardnerella vaginalis* ribonucleotide reductase (*Gv*RNR) from two isoforms: a) *Gv*R2e$_{\text{QSK-Tyr150}}$ and b) $_{\text{Gv}}$R2e$_{\text{QSK-DOPA150}}$. Sequence maps of c) *Gv*R2e$_{\text{QSK-Tyr150}}$ and d) *Gv*R2e$_{\text{QSK-DOPA150}}$ generated based on UVPD + PTCR of the 43+ charge state. Tyrosine 150 is shaded in gold to indicate that it represents the position where L-DOPA replaces tyrosine in the engineered protein. The sequences shown here do not contain the initiator methionine that is cleaved co-translationally which accounts for the one residue shift in all the amino acid numbering.

**Table S1:** List of all HpaB mutations.

| Name | Fold change over wildtype | Mutations | | | | | | | | | | | | |
|---|---|---|---|---|---|---|---|---|---|---|---|---|---|---|
| | | Lib 1 | | | | Off-target | | | | | | | | |
| | | G209 | S210 | A211 | Q212 | | | | | | | | | |
| GMRQ | 10 | G | M | R | Q | | | | | | | | | |
| GARL | 10 | G | A | R | L | | | | | | | | | |
| GDRY | 13 | G | D | R | Y | | | | | | | | | |
| GSKY | 11 | G | S | K | Y | | | | | | | | | |
| GLRL | 18 | G | L | R | L | | | | | | | | | |
| GGRM | 16 | G | G | R | M | | | | | | | | | |
| GSRR | 16 | G | S | R | R | | | | | | | | | |
| GMRA | 12 | G | M | R | A | | | | | | | | | |
| GTRH | 14 | G | T | R | H | | | | | | | | | |
| GDRK | 17 | G | D | R | K | | | | | | | | | |
| PLRQ | 14 | P | L | R | Q | | | | | | | | | |
| GDRK R94H | 16 | G | D | R | K | R94H | | | | | | | | |
| | | | | | | | Lib 2 | | | | | | | |
| | | | | | | | A156 | I157 | V158 | N159 | | | | |
| GDRK R94H ALVH | 25 | G | D | R | K | R94H | A | L | V | H | | | | |
| GDRK R94H ALVN | 26 | G | D | R | K | R94H | A | L | V | N | | | | |
| GDRK R94H CIVN | 24 | G | D | R | K | R94H | C | I | V | N | | | | |
| GDRK R94H SLVN | 31 | G | D | R | K | R94H | S | L | V | N | | | | |
| GLRL ALVH | 34 | G | L | R | L | | A | L | V | H | | | | |
| GLRL ALVN | 33 | G | L | R | L | | A | L | V | N | | | | |
| GLRL CIVN | 28 | G | L | R | L | | C | I | V | N | | | | |
| GLRL SLVN | 24 | G | L | R | L | | S | L | V | N | | | | |
| | | | | | | | | | | | Lib 3 | | | |
| | | | | | | | | | | | I146 | T201A | R287 | T398A |
| GDRK R94H SLVN T201A | 33 | G | D | R | K | R94H | S | L | V | N | | A | | |
| GDRK R94H SLVN T398A | 39 | G | D | R | K | R94H | S | L | V | N | | | | A |
| GLRL ALVH I146F | 30 | G | L | R | L | | A | L | V | H | F | | | |
| GLRL ALVH T201A | 36 | G | L | R | L | | A | L | V | H | | A | | |
| GLRL ALVH T398A | 38 | G | L | R | L | | A | L | V | H | | | | A |

**Table S2:** Relevant deconvoluted mass [Da] list for *Gv*R2e$_{QSK\text{-}DOPA150}$.

| Mass [Da] | Intensity | Relative abundance | Presumed identity |
|---|---|---|---|
| 42577.774 | 389910 | 3.6 | |
| 42597.8408, | 10,868,634.28 | 100.0 | **DOPA** |
| 42614.7917 | 6,091,322.70 | 56.0 | **DOPA+17 Da** |
| 42632.8025 | 2983590.22 | 27.5 | |
| 42654.7885 | 88807.22 | 0.8 | |
| 42678.8403 | 1466943.93 | 13.5 | |
| 42694.8296 | 1467967.11 | 13.5 | **DOPA+97 Da** |
| 42711.8387 | 1610139.31 | 14.8 | |
| 42729.8016 | 130619.16 | 1.2 | |
| 42755.7808 | 521.447.70 | 4.8 | |
| 42778.8202 | 253145.13 | 2.3 | |
| 42793.7939 | 105690.67 | 1.0 | |
| 42816.954 | 72868.62 | 0.7 | |
| 42844.8583 | 76382.51 | 0.7 | |
| 42865.8645 | 91560.05 | 0.8 | |

**Table S3:** DNA sequences of wildtype genes used in this work.

| Coding Sequences |
|---|
| HpaB |
| atgaaaccagaagatttccgcgccagtacccaacgtccgttcaccggggaagagtatctgaaa<br>agcctgcaggatggtcgcgagatctatatctatggcgagcgagtgaaagacgtcactactcat<br>ccggcatttcgtaatgcggctgcgtctgttgcccaactgtacgacgcgctacacaaaccggag<br>atgcaggactctctgtgctggaacaccgacaccggcagcggcggctatacccataaattcttc<br>cgcgtggcgaaaagtgccgacgacctgcgccagcaacgcgatgccatcgctgagtggtcacgc<br>ctgagctatggctggatgggccgtaccccagactacaaagctgctttcggttgcgcactgggc<br>gcgaatccgggcttttacggtcagttcgagcagaacgcccgtaactggtacacccgtattcag<br>gaaactggcctctactttaaccacgcgattgttaacccaccgatcgatcgtcatttgccgacc<br>gataaagtaaaagacgtttacatcaagctggaaaaagagactgacgccgggattatcgtcagc<br>ggtgcgaaagtggttgccaccaactcggcgctgactcactacaacatgattggcttcggctcg<br>gcacaagtaatgggcgaaaacccggacttcgcactgatgttcgttgcgccaatggatgccgat<br>ggcgtcaaattaatctcccgcgcctcttatgagatggtcgcgggtgctaccggctcaccgtat<br>gactacccgctctccagccgcttcgatgagaacgatgcgattctggtgatggataacgtgctg<br>atcccatgggaaaacgtgctgatctaccgcgattttgatcgctgccgtcgctggacgatggaa<br>ggcggtttcgcccgtatgtatccgctgcaagcctgtgtgcgcctggcagtgaaactcgacttc<br>attacggcactgctgaaaaaatcactcgaatgtaccggcaccctggagttccgtggtgtgcag<br>gccgatctcggtgaagtggtggcgtggcgcaacaccttctgggcattgagtgactcgatgtgt<br>tctgaagcgacgccgtgggtcaacggggcttatttaccggatcatgccgcactgcaaacctat<br>cgcgtactggcaccaatggcctacgcgaagatcaaaaacattatcgaacgcaacgttaccagt<br>ggcctgatctatctcccttccagtgcccgtgacctgaacaatccgcagatcgaccagtatctg<br>gcgaagtatgtgcgcggttcgaacggtatggatcacgtccagcgcatcaagatcctcaaactg<br>atgtgggatgccattggcagcgagtttggtggtcgtcacgaactgtatgaaatcaactactcc<br>ggtagccaggatgagattcgcctgcagtgtctgcgccaggcacaaagctccggcaatatggac<br>aagatgatggcgatggttgatcgctgcctgtcggaatacgaccagaacggctggactgtgccg<br>cacctgcacaacaacgacgatatcaacatgctggataagctgctgaaataa |
| HpaC |
| atgcaattagatgaacaacgcctgcgctttcgtgacgcaatggccagcctgtcggcagcggta<br>aatattatcaccaccgagggcgacgccggacaatgcgggattacggcaacggccgtctgctcg<br>gtcacggatacaccaccatcgctgatggtgtgcattaacgccaacagtgcgatgaacccggtt<br>tttcagggcaacggtaagttgtgcgtcaacgtcctcaaccatgagcaggaactgatggcacgc<br>cacttcgcgggcatgacaggcatggcgatggaagagcgttttagcctctcatgctggcaaaaa<br>ggtccgctggcgcagccggtgctaaaaggttcgctggccagtcttgaaggtgagatccgcgat<br>gtgcaggcaattggcacacatctggtgtatctggtggagattaaaaacatcatcctcagtgca<br>gaaggtcacggacttatctactttaaacgccgtttccatccggtgatgctggaaatggaagct<br>gcgatttaa |
| PP2551 |
| atgaatccgactttcgcaagcctgtccctggcgcacctgcgcactctggatcacctgctgcaa<br>ctgaaaaacctgagccacgcagcagaacgcctgggtgtatcccagtctgccctgtctcgtcag<br>ctggcccatctgcgtgaggcgttcgatgatccgctgctggtccgtcaaggccgtggttatgta<br>ctgtctgaacacgcagaggccctggtcgaaccgctgcgtcaggtactggaggagctgcacgct<br>ctgcgccagcctgctatcttcgatccggcccgctgcgaacgtcgcttttgcctggcagcgagc<br>gattacgtagcggagcatatgctgccgctgctggttgcggcactggagcgtgaagctccaggt<br>gtttctctggagtaccgtacctggcaggcgggccaatatgctctgctggcttctggcgagatc |

| |
|---|
| gacctggccaccactctgttcgatgagtccccgccgaacctgcacggtcgtctgctgggtgaa<br>gatcgcgctgtatgtctgatgcgtcaggatcacccgctggctgcacaggcagcgctgtctcag<br>gcagactacctggcgtacaaacacgtgcgtatcagcggtggtggtgataaagactctttcatc<br>gatcgccatctgcgcgcccagggcctgcagcgtcgcgtgtccctggaagtaccattcttctgc<br>gcgaccgtacaggtgatcgcgtcttctcaggctgttgccacggtgccagagcatattgctcgc<br>cagctgtcccgcctgcacgatctggcgtggcgcccgctgggcttcattgaccacagccagcgt<br>tactgggtggtttggcaccagcgcctgcaggcgtctgctgagcaccgttggctgcgtaaccgt<br>gttttcgaactgtggcgtcagtctcagttcggtgtacagggtggccatgcaggttctccgtaa |

**Table S4:** DNA sequences of the promoter/operator sequence with and without optimizations and an RBS used in this work. The nominal start codon is bolded. The terminator is highlighted in teal, the added insulating sequence in red, and the RBS highlighted in yellow.

| Promoter/Operator/RBS Sequences |
|---|
| Minimal PP2552 Promoter/Operator Sequence |
| catagcagctatgcggtaagcgaggttattcggctggggataggtgcctagactggggcattgtgttgattgtgcggcttc<br>ttcgcggctgtaggcgcgggtttacccgcgaaagggccagcacaggcaatggataaccc |
| Optimized PP2552 Promoter/Operator/RBS sequence |
| AACGcgacgttctccaggcatcaaataaaacgaaaggctcagtcgaaagactgggcctttcgttttatctgttgtttgtc<br>ggtgaacgctctttaccgctctcccttatgcgactccatagcagctatgcggtaagcgaggttattcggctggggataggt<br>gcctagactggggcattgtgttgattgtgcggcttcttcgcggctgtaggcgcgggtttacccgcgaaagggccagcac<br>aggcaatggataaccctaaggaggtacgtacat**ATG** |
